# Structural basis of nearest-neighbor cooperativity in the ring-shaped gene regulatory protein TRAP from protein engineering and cryo-EM

**DOI:** 10.1101/2024.05.02.592192

**Authors:** Weicheng Li, Haoyun Yang, Kye Stachowski, Andrew S. Norris, Katie Lichtenthal, Skyler Kelly, Paul Gollnick, Vicki H. Wysocki, Mark P. Foster

**Affiliations:** Department of Chemistry and Biochemistry, The Ohio State University; Center for RNA Biology, The Ohio State University; Native MS Guided Structural Biology Center, The Ohio State University; Department of Biology, University at Buffalo

## Abstract

Homotropic cooperativity is widespread in biological regulation. The homo-oligomeric ring-shaped *trp* RNA binding attenuation protein (TRAP) from bacillus binds multiple tryptophan ligands (Trp) and becomes activated to bind a specific sequence in the 5’ leader region of the *trp* operon mRNA. Ligand-activated binding to this specific RNA sequence regulates downstream biosynthesis of Trp in a feedback loop. Characterized TRAP variants form 11- or 12-mer rings and bind Trp at the interface between adjacent subunits. Various studies have shown that a pair of loops that gate each Trp binding site is flexible in the absence of the ligand and become ordered upon ligand binding. Thermodynamic measurements of Trp binding have revealed a range of cooperative behavior for different TRAP variants, even if the averaged apparent affinities for Trp have been found to be similar. Proximity between the ligand binding sites, and the ligand-coupled disorder-to-order transition has implicated nearest-neighbor interactions in cooperativity. To establish a solid basis for describing nearest-neighbor cooperativity we engineered dodecameric (12-mer) TRAP variants constructed with two subunits connected by a flexible linker (dTRAP). We mutated one of the protomers such that only every other site was competent for Trp binding. Thermodynamic and structural studies using native mass spectrometry, NMR spectroscopy, and cryo-EM provided unprecedented detail into the thermodynamic and structural basis for the observed ligand binding cooperativity. Such insights can be useful for understanding allosteric control networks and for the development of new ones with defined ligand sensitivity and regulatory control.

**Significance:** Homo-oligomeric proteins are ubiquitous in biology, and their function is often regulated by activator ligands that bind cooperatively, via poorly understood mechanisms. TRAP is a bacterial protein responsible for sensing changes in cellular tryptophan concentration, and for downregulating its production when levels are elevated. By fusing two subunits via a flexible linker, we assembled protein rings in which every other of its twelve ligand binding sites is incapable of binding. We used calorimetry and native mass spectrometry to measure ligand binding and found that these nearest-neighbor interactions are essential for cooperative ligand binding. Cryo-EM of these engineered proteins provided the means to observe the structure-thermodynamic linkage essential for understanding ligand binding cooperativity.

## Introduction

The Bacillus *trp* RNA-binding attenuation protein (TRAP) functions to control expression of the *trp* operon and therefore production of tryptophan (Trp) [1]. TRAP comprises a small ∼74 residue polypeptide that assembles into homo-oligomeric ring-shaped structures of 11 to 12 subunits, depending on species, that bind free Trp in sites located at the interface between adjacent subunits. Upon binding to multiple Trp ligands, TRAP is activated to bind with high affinity to a specific sequence containing multiple triplet repeats of GAG and UAG located in the 5’-leader region of the *trp* operon mRNA. These triplet repeats overlap stable secondary structural elements, and TRAP binding destabilizes and/or prevents their formation. The RNA structures that are formed upon TRAP binding play roles in down-regulating expression of the Trp biosynthesis machinery by promoting both transcription attenuation and translational repression [1]. Because of these properties TRAP has served as a paradigm for understanding ligand-modulated gene expression [1–3], ligand-coupled protein folding [4,5], protein-mediated RNA remodeling [6–8], and homotropic and heterotropic cooperativity [9–16], and as a tool for nanotechnology [17,18]. As a highly specific ligand-activated gene regulatory protein [12,19–21], TRAP also has potential as a template for systems biology applications [22].

Trp regulates TRAP by inducing local structural changes in protein loops spanning their binding sites [4,5,9,15,23]. Crystal structures have been determined for TRAP from *Bacillus subtilis* (*Bsu*)*, Geobacillus stearothermophilus* (*Gst*, previously *B. stearothermophilus* [24]) and from *Alkalihalobacillus halodurans* (*Aha,* previously *Bacillus halodurans* [24]). These structures, along with data from native mass spectrometry (nMS) [11,16,25,26], established that TRAP proteins assemble into predominantly undecameric (11-mer) or dodecameric (12-mer) rings featuring β-sheet structures that extend from one protomer to the next. The *Bsu* and *Gst* proteins are primarily undecameric, while the *Aha* forms dodecameric rings (Figure 1). The Trp ligands bind in the protomer-protomer interface in sites that are gated by loops (BC and DE) shown to be flexible in the absence of Trp [4,5,27]. In the presence of Trp, the flexibility of these loops is attenuated, and RNA binding on the perimeter of the ring is facilitated by successive contacts to these rigidified regions of the TRAP rings.

**Figure 1.**
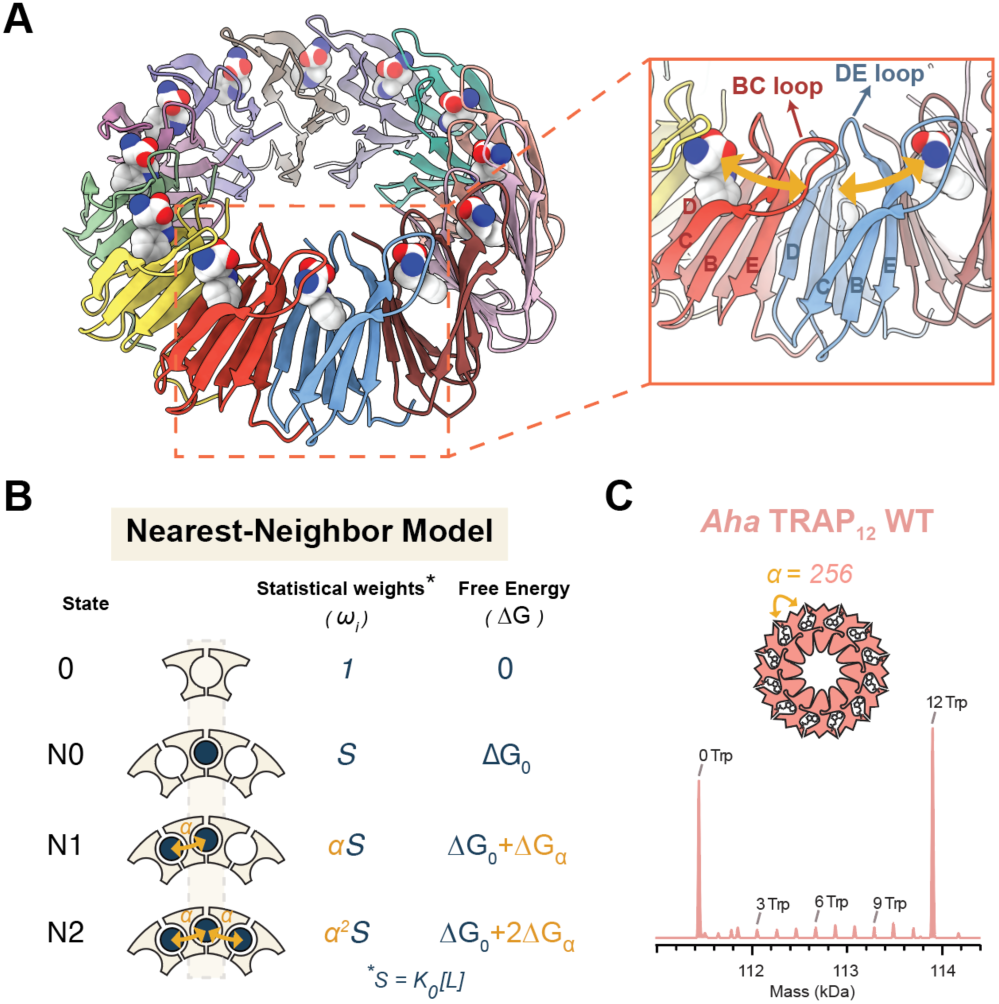
Dodecameric TRAP from Alkalihalobacillus halodurans (Aha) exhibits strong nearest-neighbor (NN) cooperativity in Trp binding. (A) Crystal structure of the ligand activated Aha TRAP 12-mer (PDB:3ZZL) shows 12 bound Trp ligands (spheres) located between adjacent protomers (cartoon). Adjacent Trp binding sites are separated by ∼12 Å via a seven-strand β-sheet that connects the protomers. Inset: Loops BC and DE that enclose the Trp binding site of one protomer connect it to adjacent Trp binding sites. (B) Configurations and parameters of the nearest-neighbor (NN) statistical thermodynamic model applied to TRAP. Each ligand binding site can populate one of four possible energy states: a reference empty state (state 0), and states bound with no occupied neighbors (state N0), bound with a single occupied neighbor (state N1), or bound with two occupied neighbors (state N2). Statistical weights ω are assigned relative to the reference state 0, with S=K_0_[L], where K_0_ = exp(-ΔG_0_/RT) is the equilibrium association constant, [L] is the free ligand concentration and α = exp(-ΔG_α_/RT) is the NN interaction term that relates the affinity of a site with an occupied neighbor to an isolated site (α = K_N1_/K_N0_). (C) (Top) The NN interaction term α defines thermodynamic coupling between adjacent Trp binding sites in Bha TRAP. (Bottom) Deconvolved native mass spectrum of a mixture of 60 µM Bha TRAP sites with 40 µM Trp; fitting a series of such spectra using the NN model revealed an interaction term α=256, corresponding to nearest-neighbor interaction of -3.3 kcal/mol ^1^.

Structural features of the Trp binding sites in TRAP rings are suggestive of homotropic Trp-Trp binding cooperativity. The BC and DE loops that gate access by each Trp ligand to an individual site also structurally connect each binding site to its neighbor (Figure 1A). In particular, the BC loop in *Aha* TRAP features several residues that form polar contacts to the bound Trp (Thr30 Oγ, Thr25 O, Gly27 O and N) [28], and also connects two Trp binding sites via residues in β strands B and C (His33, His34, Gly23, Val21). Similar site-site connections are observed in the structures of the *Bsu* and *Gst* proteins (e.g., as seen in 1WAP, 1QAW). Because of this arrangement, binding of Trp to one site has the potential to affect the structure and thermodynamics of Trp binding to adjacent sites, resulting in thermodynamically inequivalent binding to otherwise identical sites.

Although homotropic cooperativity is a common feature of homo-oligomeric ligand binding proteins [29], deciphering the structural basis for cooperative behavior remains a major challenge. One of these challenges has to do with measuring the energy transduction between sites – that is, to establish how much the free energy of binding is altered by other ligand binding events. Part of this difficulty arises from the fact that ligand binding events in oligomeric proteins follow statistical, not sequential pathways, so that observed binding is a population weighted average of weak and strong binding events. A second challenge is in linking those thermodynamic quantities to changes in the structure or conformational landscape sampled by the protein.

For molecules in which a mechanism of cooperativity can be reasonably inferred from the overall structure, the thermodynamic part of the puzzle can be simplified by formulating an appropriate partition function that parametrizes coupling energy between sites [10,30,31]. In the case of homo-oligomeric ring-shaped proteins, a plausible mechanistic hypothesis is that cooperativity between ligand binding sites arises from nearest-neighbor (NN) interactions (Figure 1B) [10,11,16,31]. A binding polynomial built on such an Ising-like lattice has been formulated for *Gst* and *Aha* TRAP, and demonstrated to explain and reproduce ligand binding thermodynamic data [11,16]. For *Gst* TRAP, fitting of Trp-TRAP binding data from isothermal titration calorimetry (ITC) and nMS, found a modest favorable coupling energy of −1.2 kcal/mol was obtained, whereas for *Aha* TRAP, a much larger coupling energy of −3.3 kcal/mol at 25°C was required to explain the observed binding isotherms [16]. These coupling free energies correspond to cooperativity factors α of 8 and 256, describing the *fold-change* in the intrinsic binding equilibrium constant *K*_0_ (Figure 1B).

With these thermodynamic parameters in hand, it remains challenging to determine the structural basis for this thermodynamic coupling because high-resolution structural methods are applied to ensembles containing statistical mixtures of states. That is, for a population of molecules in which, on average half of the sites are occupied ligands, there are 2*^n^* possible configurations with 0 to *n* bound ligands, and the observed structural features will be averaged among this population. For a dodecameric protein, there are 2^12^ = 4096 possible states! In the absence of cooperativity, these 2*^n^* states will follow a binomial pattern, whereas positive cooperativity has the effect of depleting the population of partially bound states, favoring the fully bound and empty states. At the extreme of very high cooperativity, the fully bound and empty states will dominate the distribution (e.g., Figure 1C), and structural signals will be dominated by those species. That makes it impossible to discern the effect of one ligand on the structure of other sites in the same lattice.

To determine the structural basis of Trp-Trp cooperativity in TRAP, it would be helpful to be able to assemble and determine the structure of a homogeneous population of rings in which *every-other*-*site* is occupied by Trp. Then, by comparison to the structure of the ligand-free *apo* and ligand-bound *holo* states, the effect of Trp binding on the adjacent sites could be determined.

Such an arrangement is not possible with undecameric TRAP, since 11 is a prime number and preparation of such a homogeneous sample with alternating sites occupied necessarily requires even number symmetry. However, by linking together two protomers, destroying one of the binding sites, and then assembling “dodecamers” with these dimeric subunits, such arrangements could in principle be achieved. Such samples would also serve as an important test of the validity of a postulated NN cooperativity model, as destroying every other site should result in a loss of cooperative behavior.

Informed by prior experiments with *Gst* and *Aha* TRAP [26,28,32], we engineered a series of protein constructs in which two protomers of *Aha* TRAP are fused via a flexible linker (termed here dTRAP; Figure 2). nMS confirms the stoichiometries of the assembled oligomers while solution NMR spectroscopy indicates that the linker minimally perturbs the structure of the protein. Binding assays by calorimetry and spectroscopic means similarly indicate that the linker is not strongly perturbing in the context of the wild-type protein, while mutation of one of the two binding sites results in a strong binding defect, consistent with NN interactions playing a dominant role. Finally, we applied cryo-EM methods and single particle reconstruction methods on dTRAP in its ligand-free *apo* state, its ligand-bound *holo* state, and of the mutant (WT-Mut) with every other site occupied to obtain insights into the structural basis of the strong cooperative binding observed in *Aha* TRAP. Such insights can be useful for understanding allosteric control networks and for the development of new ones with defined ligand sensitivity and regulatory control.

**Figure 2.**
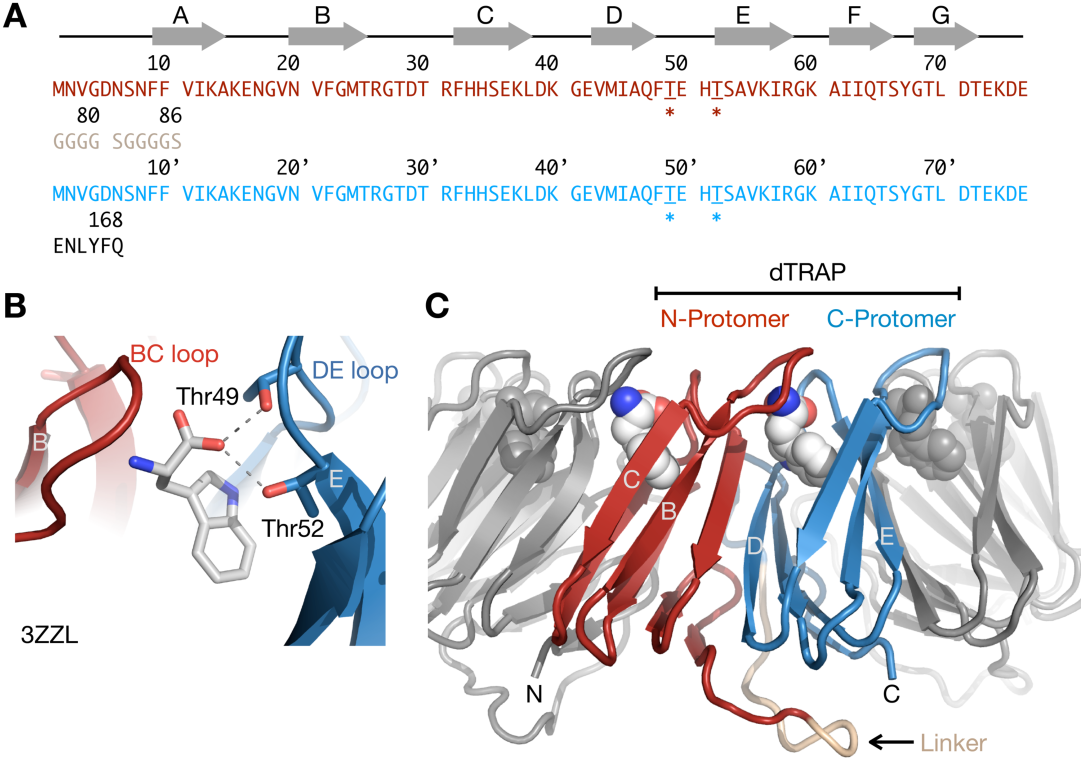
Design of linked dTRAP to enable control over nearest-neighbor configurations. (A) Primary sequence of tandem Aha dTRAP linked protomers. Secondary structural elements observed in the crystal of the wild-type protein, 3ZZL are indicated schematically. The Gly-Ser linker is grey, residual C-terminal TEV-cleavage site is black. Thr49 and Thr52, mutated to Ala to disrupt Trp binding are indicated by “*”. (B) Hydrogen bonds between the sidechain hydroxyls of Thr49 and Thr52 and the Trp ligands, as observed in the crystal, 3ZZL. (C) Rosetta model of dTRAP based on 3ZZL. The 10-redidue linker is predicted to be at the base of the TRAP ring and not interfere with Trp or RNA binding. Protomers colored as in A.

## Results and Discussion

### Engineered linked TRAP dimers (dTRAP) enable control over nearest-neighbor configurations

To manipulate the NN effect in TRAP, we engineered pairs of tandemly linked *Aha* TRAP subunits fused with flexible linkers, here termed dTRAP (Figure 2). Because Trp binds between adjacent protomers, in dTRAP one of the Trp binding sites lies at the interface connected by the linker, while the other binding site is between linked dimers. By introducing mutations to disable Trp binding on either the N- or C-terminal protomer, only every other site would be competent for Trp binding, and the potential for NN interaction would be eliminated (Figure 2B, C). In the crystal structure of *Aha* TRAP [28], the side chain hydroxyls of Thr49 and Thr52 from the DE loop of each protomer are observed to hydrogen bond with one of the carboxyl oxygens of the bound Trp ligand. Mutation to Ala of both of these residues, T49A/T52A, is expected to strongly weaken Trp binding to that site [33,34].

To avoid perturbing the structure and thermodynamics of Trp binding by TRAP we experimented with three different linker lengths for fusing C- to N-terminus two *Aha* TRAP protomers with flexible Gly-Ser linkers (GGGS, GGGSGGG and GGGGSGGGGS)[35,36]. We then evaluated the resulting fusion proteins in terms of recombinant expression yield, *in vitro* solubility, and Trp- and RNA-binding capabilities (Figure S 2). Based on those criteria, and guided in part by loop modeling with Rosetta [37] and AlphaFold2 [38] (Figure 2, Figure S 3), we selected for biochemical characterization a fusion construct with the 10-residue linker (Gly_4_Ser)_2_ .

### *Aha* dTRAP retains native oligomeric structure and strong cooperativity in Trp binding

By several measures, linking together two WT protomers with the (G_4_S)_2_ linker did not significantly disrupt the structural integrity of the protein ring, or its ligand binding capacity. Two-dimensional ^1^H-^15^N-correlated NMR spectra were recorded for both the ligand-free *apo* and ligand-saturated *holo* states of both *Aha* TRAP and dTRAP (Figure 3A). *Aha* TRAP protomers comprise 76 amino acids and both *apo* and *holo* spectra feature the number of signals expected for a single chain, indicating that the protein exhibits C12 cyclic symmetry. Although additional signals are expected from the linker and C-terminal protease site in dTRAP (Figure 2), most signals superposed well in both *apo* and *holo* states, indicating that introduction of the linker does not significantly perturb the structure of the protein. Similarity of the Trp-dependent circular dichroism (CD) spectra indicate that the linked protein dTRAP binds Trp similarly to the WT protein, with similar *apparent* dissociation constants (Figure 3B). Isothermal titration calorimetry (ITC) data also showed that these two proteins share similar bi-phasic trends, indicative of cooperative binding (Figure S 4).

**Figure 3.**
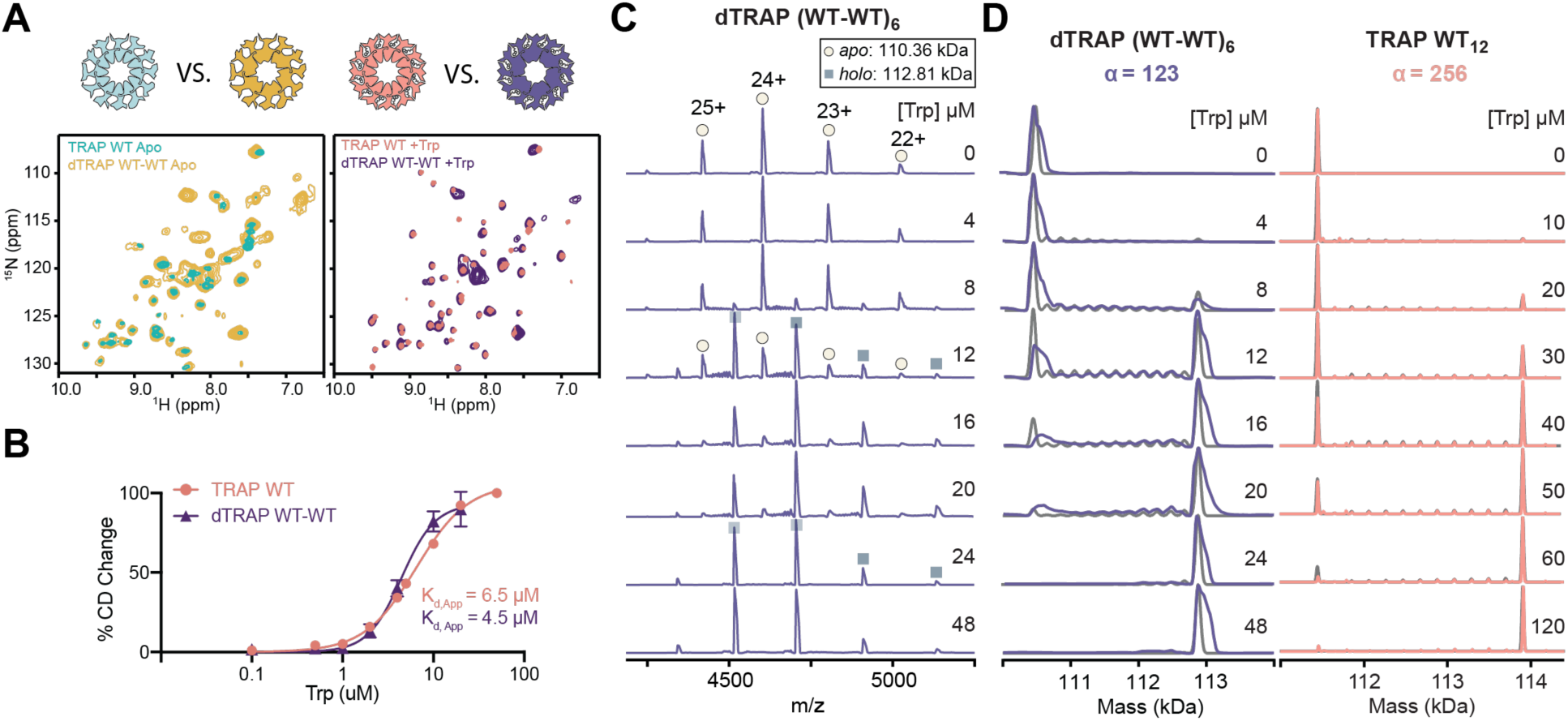
Linked dTRAP retains native structure and cooperative ligand binding. **(A)** Comparative 2D ^1^H-^15^N TROSY-HSQC NMR spectra of TRAP WT and linked dTRAP, recorded at 600 and 800 MHz, respectively, in the absence (left) and presence (right) of saturating Trp. The general similarity in peak positions indicates that introduction of the linker has a minimal impact on the structure of the oligomeric TRAP ring, and that Trp binding is minimally perturbed. **(B)** Wild-type (WT) and dTRAP exhibit similar apparent Trp binding affinities as monitored by circular dichroism (CD) at 228 nm and fit with a two-state model. **(C)** Native mass spectra of dTRAP (1.7 µM) titrated with 0-48 µM Trp. Increased Trp concentration in each titration shifts the population from apo (Trp_0_, circle) to holo (Trp_12_, square). **(D)** Deconvolved zero-charge mass spectra over a range of Trp concentrations for TRAP (left) and dTRAP (right). The experimental spectra (color) are superimposed on simulated data (grey) derived from best-fit NN model parameters. In both spectra, the dominant populations comprise the apo-state (0 Trp) and the holo-state (12 Trp), indicating strong positive cooperativity. The best-fit NN interaction terms α = 256 and α = 123, correspond to RT ln α = 3.3 and 2.9 kJ/mol at 25 °C, respectively.^1^

We have found nMS to be particularly useful for quantifying cooperativity in TRAP [11,16] and those spectra revealed similar strong cooperativity in dTRAP compared to TRAP (Figure 3D). Native mass spectra of both proteins recorded in the absence of Trp yielded a dominant set of signals corresponding to the mass of the oligomeric *apo* proteins (TRAP: 111,449 Da expected, 111,454 observed; dTRAP: 110,355 Da expected, 110,394 observed; differences between expected and measured masses are likely the result of attempts to use cool MS tuning to avoid restructuring of the complex and associated adducting with salts and solvents, in addition to uncertainty in the specific charging species; i.e., H+, Na+, K+. Across a range of Trp concentrations, nMS signals from the *apo* and fully loaded *holo* states (dTRAP with 12 bound Trp, expected: 112,805 Da, observed: 112,832 Da; TRAP with 12 bound Trp: 113,899 Da expected, 113,903 observed) dominated both sets of spectra (Figure 3C,D). Depletion of partially loaded states of the oligomer is a defining feature of positive cooperativity [16], indicating that both TRAP and dTRAP exhibit strong positive cooperativity under those conditions. Populations of Trp_n_-TRAP_12_ states obtained from the nMS peak heights were fit with the NN binding model [10,11,16] using the standard deviation from three replicates as an estimate of precision. Best fit parameters to the NN models found for *Aha* TRAP a NN cooperativity strength α = 256 ± 87 and for dTRAP α = 123 ± 5; intrinsic affinities *K*_0_ were 720 ± 140 µM and 240 ± 90 µM, respectively.

## Eliminating NN interactions impairs Trp binding

To produce dTRAP variants incapable of NN interactions, dual T49A/T52A mutations were introduced either on the N-terminal or C-terminal protomer (Figure 2) to generate Mut-WT or WT-Mut dTRAP proteins, or at both sites to generate a Mut-Mut dTRAP variant. We expressed and purified each of these dTRAP variants and performed preliminary characterization. Each protein construct eluted from a size exclusion column with the apparent molecular weight expected of a TRAP dodecamer (here with six linked dTRAP). Heteronuclear NMR spectra of WT-WT, WT-Mut and Mut-WT dTRAP were highly congruent in the absence of Trp (Figure 4A, Figure S 5), indicating highly similar structures; poor expression and solubility of the Mut-Mut variant precluded it from NMR analysis. The 2D NMR spectra of WT-WT dTRAP recorded in the presence of excess Trp showed large chemical shift perturbations, consistent with Trp binding and induced structural changes (Figure 4A). By contrast, spectra of WT-Mut and Mut-WT dTRAP were largely unperturbed under equivalent Trp concentrations indicating a Trp binding defect (Figure 4A, Figure S 5). The Mut-WT variant failed to bind Trp as assayed by other metrics (Figure S 5) so without identifying the cause of that effect, we proceeded with thermodynamic analysis of WT-Mut dTRAP, which, when assembled into rings, should be capable of Trp binding at every other site, and the Mut-Mut construct, which lacks native Trp binding sites (Figure 4B-D).

**Figure 4.**
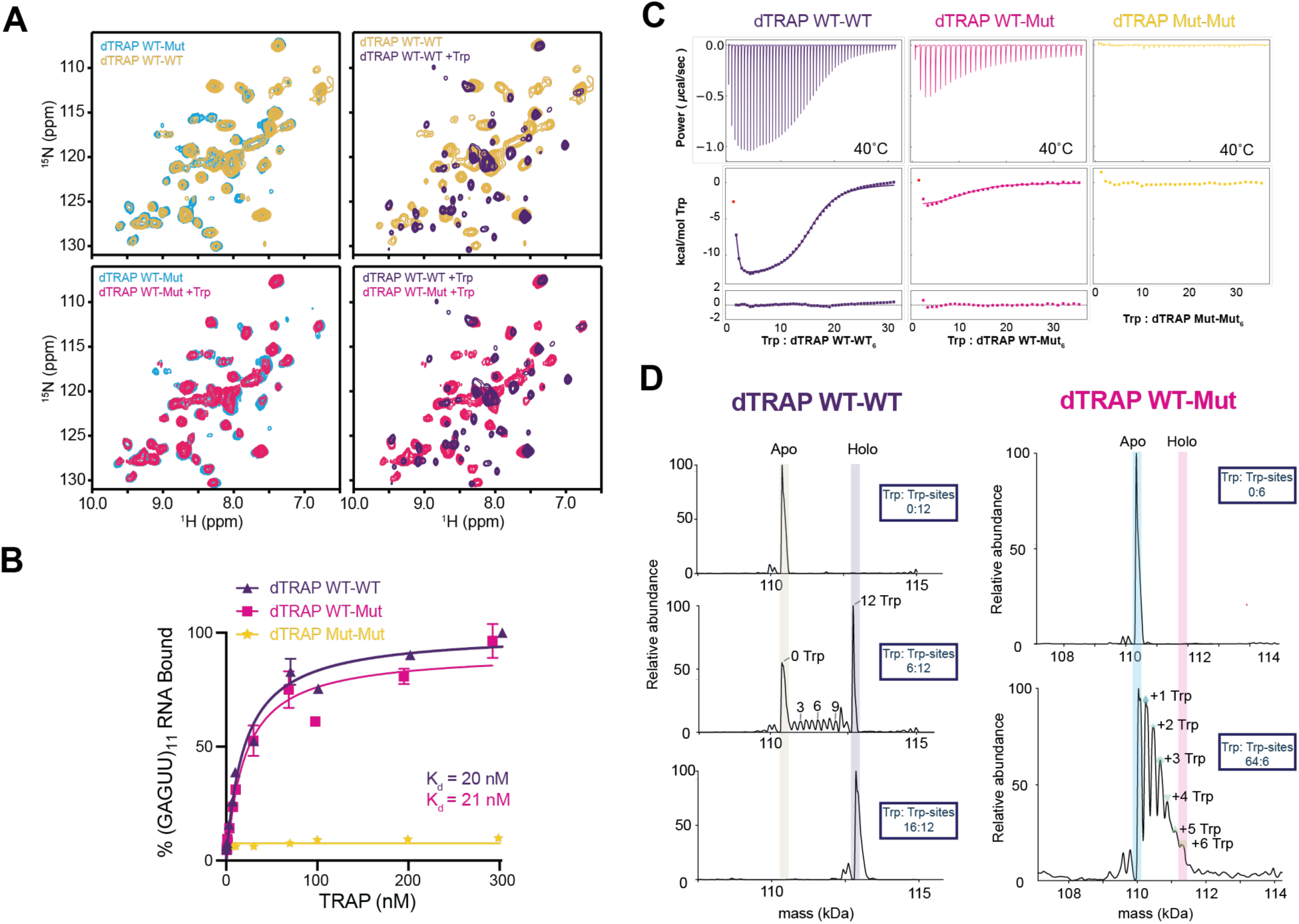
Eliminating the nearest-neighbor interaction disrupts cooperativity in dTRAP. **(A)** Comparative 2D ^1^H-^15^N-correlated NMR spectra of WT-WT and WT-Mut dTRAP in the absence and presence of Trp. Top left, WT-WT and WT-Mut adopt similar structures in the absence of Trp. Top right, WT-WT in the absence and presence of Trp. Bottom left, WT-Mut in the absence and presence of Trp; lack of perturbations reflects a binding defect. Bottom right, WT-WT and WT-Mut in the presence of Trp contrasts with their similarity in the absence. **(B)** RNA filter-binding data as a function of protein concentration for dTRAP variants towards an idealized (GAGUU)_11_ single-stranded leader RNA, in the presence of 1 mM Trp. The WT-WT and WT-Mut variants exhibit similar apparent RNA affinity, whereas the Mut-Mut variant is unable to bind RNA over the same range of TRAP concentrations. **(C)** ITC of Trp into dTRAP variants. The thermogram for WT-WT dTRAP is biphasic, indicative of cooperativity. Thermograms of dTRAP WT-Mut and Mut-Mut exhibit small and no enthalpy changes, respectively. **(D)** Deconvolved zero charge native MS spectra of Trp-dTRAP titrations (at 12 µM sites) reveal strong positive cooperative binding for WT-WT dTRAP (left) (high abundance in both apo- and holo-state), whereas dTRAP WT-Mut cannot be saturated at > 10-fold stoichiometric excess under these conditions (right).

dTRAP variants remain functional in Trp-activated RNA binding. We examined Trp-dependent RNA binding by dTRAP variants to (GAGUU)_11_ RNA repeats with a filter binding assay (Figure 4B). In the presence of 1 mM Trp, WT-Mut dTRAP maintains strong RNA binding affinity with an apparent K_d_ of 21 nM, which is close to that observed for WT-WT of 20 nM. Interestingly, although as mentioned above, Mut-WT dTRAP failed to bind Trp on its own, it does exhibit Trp-dependent RNA binding with an apparent affinity K_d_ = 27 nM (Figure S 6). These results indicate that the (G_4_S)_2_ linker is not strongly perturbing of dTRAP structure and ligand binding. In addition, the Thr/Ala mutations at every other site does not significantly diminish the protein’s ability to bind RNA. Conversely, dTRAP Mut-Mut appears incapable of RNA binding, presumably due to its inability to bind Trp (Figure 4B).

One hallmark of cooperative ligand binding is biphasic thermograms in ITC experiments [39]. ITC measurement of Trp titrations reveal a biphasic thermograms for dTRAP consistent with cooperative Trp binding and an apparent maximal enthalpy change of -13 kcal/mol (Figure 4C, Figure S 7). Under comparable binding site concentrations of WT-Mut dTRAP (∼ twice to account for WT binding sites per chain), the thermograms for titrating Trp into WT-Mut and Mut-Mut, exhibit a smaller binding ΔH of -4 kcal/mol and reduced biphasic character, whereas no measurable heat of binding was detected at that protein concentration for Mut-Mut dTRAP (Figure 4C). Although the biphasic character of the thermograms was much reduced, they were reproducibly biphasic, indicating that despite deficient Trp binding to Mut sites, WT-Mut dTRAP retains some modest cooperativity. Nevertheless, these data are consistent with a *dominant* role of NN contributions on cooperativity and overall Trp binding affinity.

Because nMS can better describe the distribution of Trp-TRAP states than can other bulk measures of ligand binding [11,16,40], we used nMS to compare Trp binding to dTRAP and WT-Mut dTRAP (Figure 4D, Figure S 8). Under conditions of approximately half maximal saturation for dTRAP at 12 µM binding sites, the mass spectrum is dominated by signals from the Trp-free *apo* protein and from the fully Trp-loaded *holo* dTRAP ring (expected: 112,805 Da, observed: 112,832 Da). By comparison, WT-Mut dTRAP exhibits a strong Trp binding defect. In the absence of Trp we observe the expected major signal corresponding to the apo WT-Mut oligomer (109,993 Da expected, 110,029 Da observed), while in the presence of a 10-fold excess of Trp (12 µM WT sites, 128 µM Trp), not only are the fully Trp-loaded rings a minor species, but intermediate species are much more populated, resembling a statistical gaussian-like distribution. Nevertheless, the population distribution of WT-Mut dTRAP rings with zero to six bound Trp was poorly fit with a non-cooperative, independent sites model, yielding non-random residuals (Figure S 9). Fitting the WT-Mut dTRAP populations using the NN cooperative model yielded small, near random residuals and best fit values of 873 ± 90 µM and 5 ± 1 for *K*_0_ and the cooperativity factor α. Thus, cooperativity in WT-Mut dTRAP is down by a factor of ∼25 compared to that of WT-WT dTRAP (α = 123). These data support a model in which the strong cooperativity observed in dTRAP arises from favorable cooperative interactions between adjacent binding sites. It also suggests that differences in the structural features of Trp-loaded oligomers composed of dTRAP and WT-Mut dTRAP may reveal insights into the basis for this cooperative effect.

## Cryo-EM single particle reconstruction of dTRAP WT-WT and WT-Mut constructs

Availability of WT-Mut dTRAP, with alternating binding-competent and -impaired sites provides an opportunity to discern the structural basis for the observed nearest-neighbor cooperativity. We sought to obtain these insights by comparing cryo-EM maps from dTRAP in the absence and presence of Trp, and of WT-Mut dTRAP in the presence of Trp; these samples are expected to exhibit characteristics corresponding to the ligand-free *apo* protein, the fully liganded *holo* protein, and an intermediate in which every other site is occupied.

Cryo-EM grids of dTRAP and its variants yielded well-defined particles with a good distribution of orientations (Figure S 10, Figure S 11, Figure S 12). Single-particle reconstructions of the Cryo-EM data with application of overall six-fold symmetry resulted in distinct electron potential maps for the three samples (Figure 5). Local resolution differences reveal a progressive increase in order from the unbound *apo* dTRAP protein, to the partially bound WT-Mut protein and finally the fully bound *holo* dTRAP. Application of six-fold symmetry is justified by the linker connecting the two protomers in dTRAP, reducing overall symmetry from twelve-fold in TRAP to six-fold in dTRAP. Nevertheless, potential map refinement using C1 symmetry resulted in maps with distinct but low overall resolution, as well as visually inequivalent maps for alternating sites in Trp-loaded WT-Mut dTRAP (Figure S 13, Figure S 14, Figure S 15); imposition of C6 symmetry following homogeneous C1 refinement increased the WT-Mut map resolution from 5.3 to 4.4 Å. Use of these C6-refined maps in particle reclassification, followed by two more rounds of homogenous and non-homogeneous refinement resulted in a map with a resolution of 4.1 Å for WT-Mut dTRAP. Similar workflows for *apo* and *holo* dTRAP yielded maps with a global resolution of 4.2 Å and 3.6 Å, respectively (gold standard FSC cutoff of 0.143; Figure 5, Figure S 10, Figure S 11, Figure S 12, Table S1). The ring shape of dTRAP oligomers is clearly established by the density maps, and the highest local resolution is obtained for the β-strands forming the base and the core of each ring. The lowest local resolution is observed for regions that correspond to the Trp binding loops. No density could be assigned to the flexible linker connecting the two TRAP protomers.

**Figure 5.**
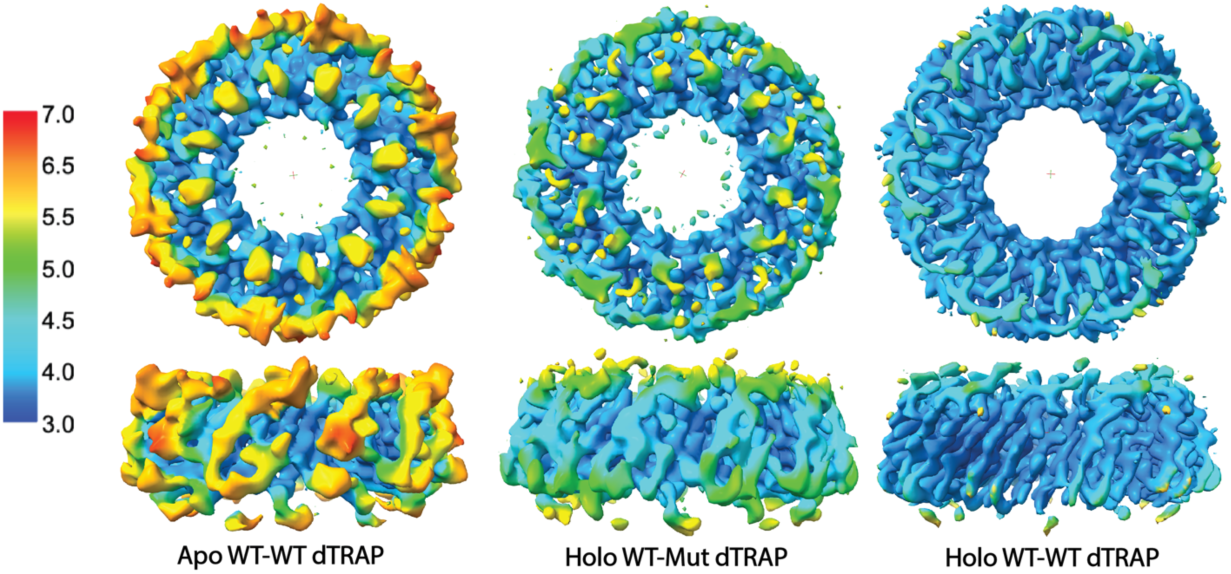
Cryo-EM of dTRAP and WT-Mut dTRAP reveal ligand-dependent ordering. Electron potential maps from single-particle reconstruction of cryo-EM images from Trp-free apo-dTRAP (EMD-44472), Trp-loaded holo WT-Mut (EMD-44473), and Trp-loaded holo dTRAP (EMD-44468) colored by local resolution, from 3 to 7 Å using a rainbow heatmap. Top, axial view of the density maps; bottom, side view. Left, apo-dTRAP, average resolution 4.2 Å, exhibits lowest resolution on the top and edges of the ring. Right, Trp-bound dTRAP, average resolution 3.6 Å, exhibits near-atomic resolution maps across the toroidal structure. Middle, Trp-bound WT-Mut dTRAP, average resolution 4.1Å, exhibits intermediate resolution. Map contour levels for apo WT, holo WT-Mut and holo WT were set in ChimeraX (www.cgl.ucsf.edu/chimerax) to 0.135, 0.11 and 0.184, respectively. Maps have been deposited to the EMBD.

The C6-refined potential maps are consistent with ligand-dependent local ordering. For the *holo* protein, maps corresponding to adjacent binding sites were highly congruent despite C6 refinement (Figure 5, Figure 6, Figure S 12). This finding is consistent with the similarity in the NMR spectra of Trp-loaded TRAP and dTRAP (Figure 3A), which indicates that the Trp bound sites are isomorphous in the Trp bound state. Similarity between the NMR spectra of TRAP and dTRAP in the absence of Trp (Figure 3A) also suggests that sites in *apo* dTRAP should be congruent. However, maps for adjacent protomers are visually different at a contour level of 0.135. This apparent asymmetry could be a consequence of the lower resolution of the *apo* maps, particularly given that the loops are expected to be dynamically disordered [4,27,41], due to the effect of the linker on particle alignment, or because the inter-protomer linker has a measurable effect on the proximal Trp binding sites. The lower resolution of the map for *apo* dTRAP, and unexpected asymmetry, gives us less confidence in interpreting this asymmetry. The electron potential map for Trp-bound WT-Mut dTRAP exhibits intermediate resolution between that observed for *apo* and *holo* dTRAP and the greatest structural difference between adjacent binding sites (Figure 5, Figure 6).

**Figure 6.**
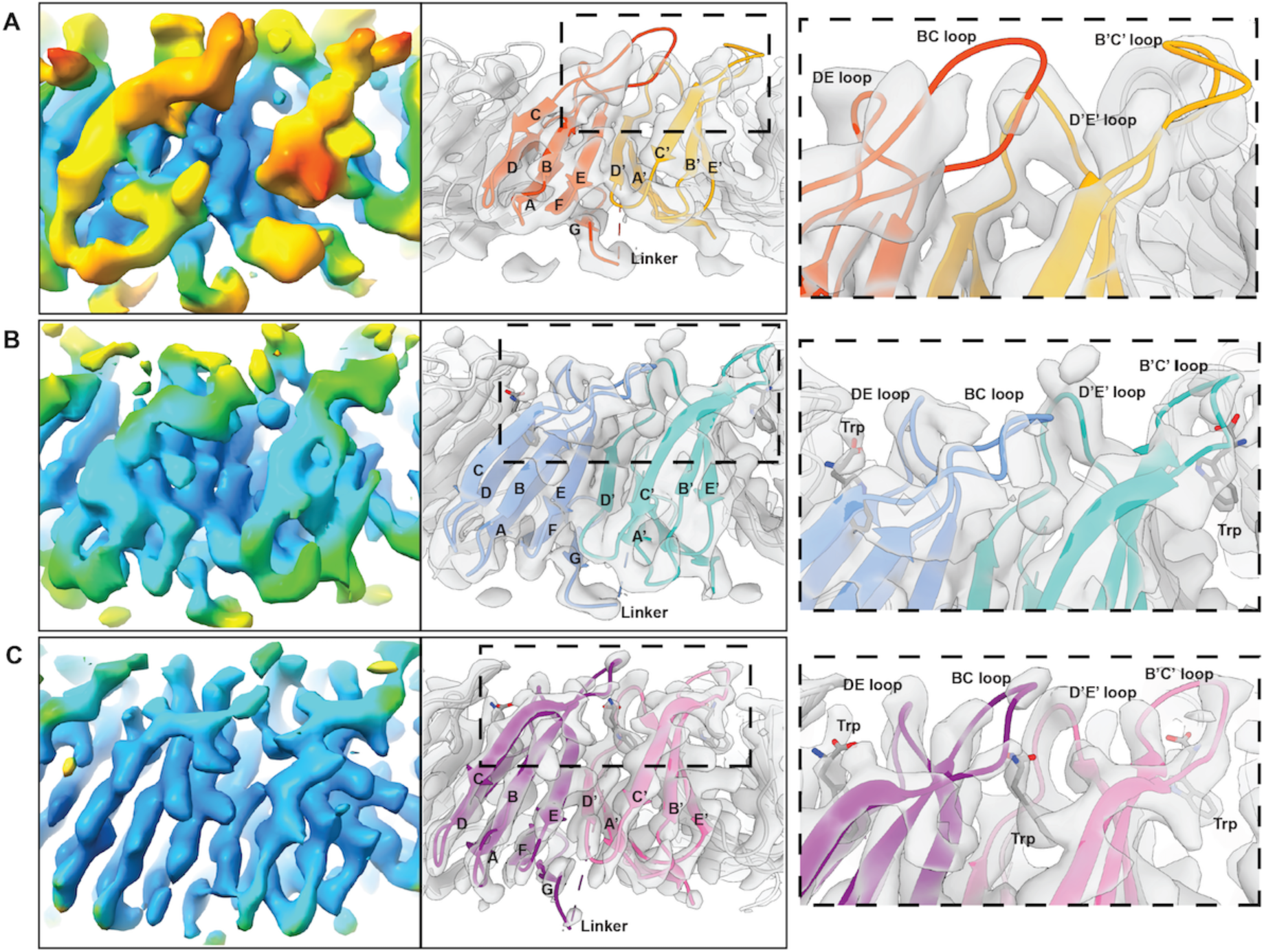
Ligand-dependent density in dTRAP linked protomer maps reveal loop stabilization. Left: Views of apo dTRAP (A), holo WT-Mut (B), and holo dTRAP (C) maps of a linked protomer colored by local resolution. Middle: models of dTRAP variants fitted into the maps. The two linked protomers are colored differently. Right: Close-up views of the BC and DE loops. Density corresponding to the BC and DE loops is mostly missing in apo dTRAP, partially discernable for the DE loops in Trp-bound WT-mutant, and much better defined in holo dTRAP. Alternating Trp-bound WT and Trp-free Mut sites has discernably different densities in WT-Mut dTRAP (B). Models have been deposited in the PDB, accession 9be7, 9be8 and 9bds, respectively.

## dTRAP rings are slightly expanded compared to *Aha* TRAP dodecamers

Comparison of crystallographic and cryo-EM data revealed expansion of the ring diameter by 6% in dTRAP compared to WT *Aha* TRAP. To build atomic models into the potential maps for *apo* and *holo* dTRAP, and Trp-bound WT-Mut dTRAP, we used Rosetta [42] to build the (Gly_4_Ser)_2_ linkers using the crystal structure of Trp-bound *Aha* TRAP (3zzl; [28]) as the template. Model refinement performed in an iterative fashion using Coot, ISOLDE and PHENIX [43–46] yielded good local agreement between the template and optimized model (Figure S 16). However, the crystallographic model of *holo Aha* TRAP unexpectedly did not fit well in the potential map of *holo* dTRAP (Figure S 17). Comparison of the crystal structure coordinates and optimized model for *holo* dTRAP revealed a 6% expansion in the diameter of the TRAP ring, as measured by the change in distance between corresponding atoms on opposite sides of the ring (e.g., Lys37 C^α^). Because the template was obtained from crystals of *Aha* TRAP, we also compared the cryo-EM electron potential map of *holo* dTRAP with that of a point mutant of *Aha* TRAP modified to form protein cages via interaction with gold nanoparticles [18]. The potential map obtained for dTRAP with its twelve effective protomers (six dTRAP chains) appears expanded by ∼6% compared to that of *Aha* TRAP dodecamers (Figure S 18). Based on this observation we conclude that the ten-residue flexible linker is the likely cause of this modest ring expansion.

## Map differences correspond primarily to the Trp binding loops

Model building into the electron potential map reveals that the largest differences in between the *apo, holo* and WT-Mut dTRAP structures are in degree of order of the BC and DE loops (Figure 6). Of the three samples, the map for *holo* dTRAP was best defined (Figure 5, Figure 6).

Backbone density could be resolved for most of the protein chain, with the exception of residues 1-6, which were also not observed in the crystal structure of wild-type *Aha* TRAP [28], the Gly-Ser linker, and the C-terminal TEV sequence; density was also weaker for Gly27 and Asp29 in the BC loop. Density corresponding to the bound Trp can be clearly observed between each protomer monomers (Figure 6C). As noted above, compared to the crystal structure of Trp-bound *Aha* TRAP, the ring is slightly expanded. Otherwise, there is strong agreement between placement of secondary structural elements and overall geometry of the Trp binding site, although sidechain rotamers cannot clearly be distinguished at this resolution.

In comparison to *holo* dTRAP, in addition to lower overall resolution for *apo* dTRAP, the most obvious feature is that the density corresponding to the BC and B’C’ loops of apo dTRAP (Figure 6A, middle and right panels) cannot be discerned. This is consistent with the loops being flexible and disordered in the absence of the ligand. The Trp binding sites in *apo* dTRAP feature only weak density, consistent with the sites being empty (Figure 6A, right panel). Although no clear density could be discerned for the BC and DE loops, they are nevertheless included in the model for guidance, but their position should not be interpreted as defined by the data.

The density map for Trp-bound WT-Mut dTRAP showed better overall resolution than that of *apo* dTRAP (Figure 5, 6B), with clearer density for the BC (and B’C’) loops. Defined density assignable to the Trp ligand is observed at alternating binding sites (Figure 6B middle and right panels). However, lesser density is also observed in the Mut sites, which could result from either buffer atoms or sidechain rearrangement, or partial misalignment of particles during reconstruction; similar weak density is observed in the apo WT-WT dTRAP ligand binding sites, suggesting the latter (Figure 6A). The higher local resolution indicates a higher degree of order in the Trp binding loops.

### Ligand binding induces increased order at neighboring sites

The principal objective of the present cryo-EM studies is to provide insights into the structural basis for ligand binding cooperativity by TRAP. The ligand binding experiments described above demonstrate that WT-Mut dTRAP is strongly deficient in Trp binding, in a way that is consistent with a major role for NN interactions in mediating this effect. One possible mechanism for achieving NN cooperativity would be to induce in neighboring sites a discrete structure that favors ligand binding. However, the BC and DE loops in the WT-Mut structure are much less well ordered than those of the holo protein, indicating that the protein with ligand bound at alternating sites retains a significant degree of disorder. Nevertheless, the loop disorder is measurably decreased in the Trp-bound WT-Mut protein, compared to the *apo* protein (Figure 6A,B). This suggests that NN cooperativity can be explained by a population shift from the wide range of conformations sampled by *apo* TRAP, to a narrower distribution of states that comparatively favor ligand binding. While this is evident in terms of increased order at the backbone level, it likely also arises from a narrowing of the conformational space sampled by protein side chains in the Trp binding site, and connecting one site to its adjacent sites via strands B,C.

## Conclusions

Trp binding to linked dTRAP approximates independent site binding. We engineered dTRAP to dissect the intrinsic affinity of Trp for a site, *K*_0_, from the NN interaction term α that results in higher affinity for binding next to occupied sites (Figure 1, Figure 2). Comparison of NMR spectra for *Aha* TRAP and linked dTRAP, and macroscopic binding metrics (Figure 3) suggested that the linker introduced only minor perturbations in the structure of the TRAP dodecameric ring and its Trp binding thermodynamics (i.e., a two-fold reduction in the interaction term α).

Mutation to Ala of two ligand-coordinating Thr residues in one of the protomers of dTRAP resulted in protein rings with six intact Trp binding sites, but a major defect in Trp binding, with a fifteen-fold reduction in α (Figure 4). This is consistent with a dominant effect from NN interactions between sites. Comparing cryo-EM density maps for *apo* dTRAP, WT-Mut dTRAP and *holo* dTRAP we observe progressive increase in map resolution for the BC and DE loops that gate access to the Trp binding sites (Figure 6, Figure 6).

A plausible mechanism for cooperativity in *Aha* TRAP, consistent with the presented data, is that binding of Trp to a site rigidifies the BC and DE loops flanking the site, while their proximity and structural coupling to adjacent sites (Figure 1) likewise results in partial ordering of the adjacent sites. The accompanying shift in the free energy landscape can thus be understood to favor the ligand-bound state, without specifying specifically which conformational states are excluded by rigidification of the adjacent sites (Figure 7). In this model ligand binding to neighboring sites changes the free energy landscape such that binding-competent confomrations are more populated and thus subsequent ligand binding is favored. A lesser effect from binding at next-nearest-neighbors can also be envisioned as arising from more modest shift in the free energy landscape.

**Figure 7.**
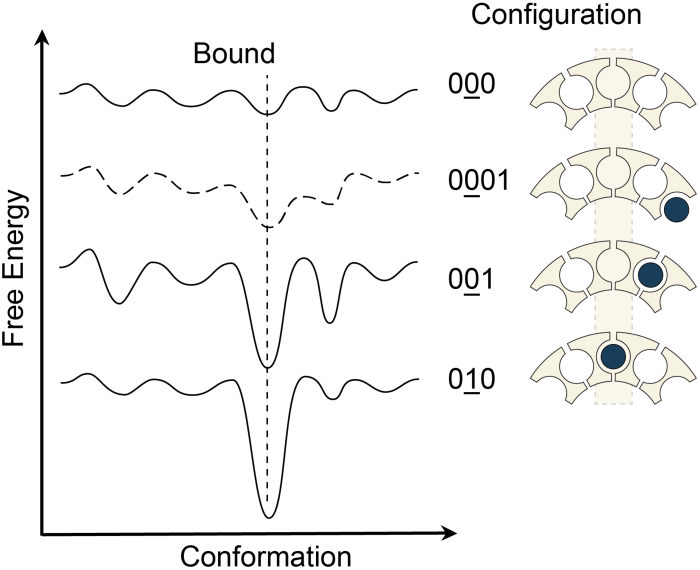
Cooperativity through population shifts induced by nearest-neighbor binding in TRAP. x-axis, idealized one-dimensional conformational coordinate; y-axis, free energy; lower energy conformations (states) are more populated. In the absence of Trp (0<u>0</u>0, top), the BC and DE loops sample a wide range of iso-energetic conformations, most of which do not favor Trp binding. The conformational landscape of a Trp-bound site (0<u>1</u>0, bottom) is narrow and deep, populating conformations corresponding to the Trp-bound state. Binding of a ligand to a neighbor (0<u>0</u>1) alters the free energy landscape of a site, favoring the bound state, but also populating other states, such as those that allow binding. Next-nearest-neighbor binding (0<u>0</u>01, dashed line) may also result in a population shift, exerting a smaller magnitude cooperative effect. Free energy curves are offset for clarity.

The combination of protein engineering with nMS and cryo-EM single-particle reconstruction provided the means to dissect direct and indirect effects of ligand binding on TRAP. nMS provides unmatched detail into the thermodynamics of Trp binding by enabling quantification of states with differing numbers of bound Trp. Construction of the dTRAP linked variants allows for control over the Trp-TRAP configurations, whereas oligomers with isolated chains would necessarily populate statistical distribution of states, some with bound neighbors, some not. The resulting well-defined configurations provided an opportunity to detail the structural differences between TRAP rings free, fully bound to Trp, and with alternating sites bound. The results strongly support a model in which cooperativity in TRAP is dominated by ligand-mediated ordering of loops that gate Trp binding to individual sites and serve as bridges connecting sites.

While the results of the present study strongly support this model, there were complications that limited the precision of the resulting insights. Although introduction of the linkers seemed, based on NMR data and bulk measurements to not perturb the structure and thermodynamics of Trp binding (Figure 3), the linkers had a measurable effect. nMS of Trp binding to WT dTRAP nevertheless showed that ligand binding thermodynamics was slightly perturbed (by -*RT* ln Δα = -0.69 *RT*; Figure 3), and the Mut-WT protein seemed unable to bind Trp at all (Figure S 5). Moreover, the cryoEM data showed that dTRAP rings were expanded by ∼6% relative to the WT rings (Figure S 18), indicating that the linkers had a perturbing effect on the structure. In addition, while the flexibility of the linkers was intended to prevent perturbation of the protein fold and thermodynamics, the fact that the linkers are small and disordered rendered them invisible in reconstructed electron density maps. Because of this, it is difficult to assess the extent to which particle misalignment might have contributed to reduced density of the maps, especially for the Trp-loaded WT-Mut protein (Figure 5). Lastly, while the data presented strongly support a model in which disorder-order transitions are responsible for the observed Trp-Trp cooperativity, they are insufficient to map out at atomic resolution the conformational ensemble of states present in the Trp-bound and partially-loaded TRAP rings.

Such atomic level insights are likely required to enable rational efforts to modulate the thermodynamic response of TRAP and molecules like it for use in diagnostics or synthetic biology applications.

## Methods

### Preparation of linked *Aha* TRAP variants (dTRAP)

DNA constructs encoding *Aha* TRAP-(linker)-TRAP-(TEV cleavage site)-His_6_ with varying length Gly/Ser linkers were obtained commercially (GenScript, Inc) in a pET21a (Novagen) plasmid; linker sequences were GGGS, GGGSGGG, and GGGGSGGGGS. Proteins were expressed in *Escherichia coli* BL21(DE3) cells and inducted by adding 0.5 mM IPTG (isopropyl ß-D-1-thiogalactopyranosid) at mid-log phase (OD600 = 0.4 to 0.6) at 37°C. Cells were then harvested through centrifuge and then resuspended in loading buffer (20 mM NaPi, 500 mM NaCl, 100 mM Imidazole, pH = 7.4). The cells were lysed using French press method at 10,000 psi. The lysate was cleared by centrifugation at 30,000g for 20 min, and the supernatant was filtrated through 0.45 um cellulose acetate filters (Advantec) using 30mL syringes (BD). The supernatant was loaded onto 1 ml HisTrap HP column (GE Healthcare) on an AKTA fast protein liquid chromatography (FPLC). Proteins were then eluted with a gradient of imidazole from 100 mM to 1 M in the buffer of 20 mM NaPi pH 7.4, and 500mM NaCl. The C-terminal 6xHis-tag was then removed by treatment with TEV protease (in-house produced) at a protease-to-target molar ratio of 1:100 and products were passed through Ni-NTA to remove uncleaved protein. To ensure complete removal of bound Trp, proteins were dialyzed in 3.5 kDa cut-off (Spectrum Labs) dialysis tubing against 3 M Gdn-HCl in buffer A (100 mM NaCl, 50 mM NaPO_4_, pH 8.0 at 25 °C, 0.02% NaN_3_) for 6 hours at room temperature. The samples were then step-down dialyzed against 1.5 M and 0 M Gdn-HCl in buffer A. Refolded proteins were further purified by size-exclusion chromatography using a Hiload 16/600 Superdex 75 column (Sigma-Aldrich) in buffer A.

After initial trials for expression, solubility, and Trp- and RNA-binding activity, the construct with the GGGGSGGGGS linker (G_4_S)_2_, was selected for further characterization. The resulting protein has the expected amino acid sequence: MNVGDNSNFF VIKAKENGVN VFGMTRGTDT RFHHSEKLDK GEVMIAQF**T**E H**T**SAVKIRGK AIIQTSYGTL DTEKDE<u>GGGG</u> <u>SGGGGS</u>MNVG DNSNFFVIKA KENGVNVFGM TRGTDTRFHH SEKLDKGEVM IAQF**T**EH**T**SA VKIRGKAIIQ TSYGTLDTEK DEenlyfq, where the linker is underlined, and the C-terminal TEV protease recognition site is lowercase. With a predicted molecular weight (MW) of 18,392.45 Da (ProtParam), fully assembled Trp-free rings have an expected MW of 110,355 Da.

To produce dTRAP variants with one or both sites deficient in Trp binding, we used site-directed mutagenesis to generate Ala substitutions of Thr49 and Thr52 (boldface, above) in the N-terminal protomer, the C-terminal protomer, or both protomers to generate dTRAP WT-Mut, dTRAP Mut-WT, and dTRAP Mut-Mut proteins, respectively. Sample preparation steps was as above, except that dialysis in 1 M Gdn-HCl buffer A was enough to deplete the bound Trp. Variants with one pair of Thr/Ala mutants (WT-Mut, Mut-WT) have a predicted MW of 18,332.40 Da, and the double mutant (Mut-Mut) MW is 18,272.35 Da (ProtParam).

### Concentration of Components

All dTRAP proteins shared the same predicted extinction coefficient at 280 nm (ε = 4470 M^-1^cm^-1^; ProtParam). Protein concentrations were measured using a Nanodrop 2000c spectrophotometer (Thermo Scientific) in droplet mode. Unless otherwise specified, TRAP concentrations refer to the concentration of the TRAP 12-mer ring, not the protomer concentration. Ring concentration is 1/12th the promoter concentration for WT TRAP; it’s 1/6th the protomer concentration for dTRAP and its variants. The binding site concentration equals the promoter concentration for WT TRAP, it’s twice the concentration for WT-WT dTRAP, and it equals the promoter concentration for Mut-WT and WT-Mut dTRAP. Trp concentration was determined by UV absorbance at 280 nm with the extinction coefficient ε =5540 M^-1^ cm^-1^ using a Nanodrop 2000c spectrophotometer (Thermo Scientific) in droplet mode.

### Circular Dichroism (CD)

Trp binding to dTRAP variants was assayed by CD spectropolarimetry using a JASCO model J-715 instrument and a 1 mm pathlength at 37 °C. The data were recorded between 226 nm to 230 nm using 2 µM TRAP in 50 µM sodium phosphate solution (pH = 8.0) with the Trp concentrations ranging from 0 µM to 300 µM.[16,47] The CD signal at 228 nm was used to quantify tryptophan binding [13,23]. The fractional change in CD signal *Y* was fit with the single-site mass-balance binding model, with parameters describing total protein and ligand concentrations, [*P_tot_*], [*L_tot_*], and apparent dissociation constant, *K*_D_:

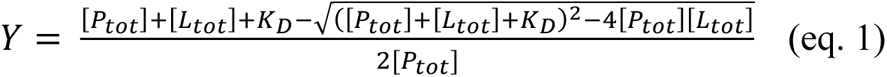

### Isothermal Titration Calorimetry (ITC)

ITC isotherms were recorded on the MicroCal VP-ITC instrument (Malvern) at temperatures of 35 and 40 °C. HPLC-purified, refolded, and SEC S75 column-purified TRAP was dialyzed against buffer A (100 mM NaCl, 50 mM NaPO_4_, pH 8.0, 0.02% NaN_3_). Tryptophan (Trp) stock solution was prepared by dissolving L-Trp powder (Sigma-Aldrich) in the TRAP-dialysate. At 35°C, 293 µM Trp was titrated into 1.9 µM WT TRAP; 512 µM Trp was titrated into 3.4 µM dTRAP WT-WT. At 40°C, ITC thermograms of Trp into dTRAP variants were recorded at 40°C at concentrations of 435 µM Trp into 3.4 µM WT-WT, 500 µM Trp to 8.3 µM WT-Mut and Mut-WT, and 530 µM Trp to 8.3 µM Mut-Mut. Baseline correction and integration of ITC thermograms were performed using NITPIC.[48] Processed data were then fit with one-site phenomenological binding model via *itcsimlib.*^6^

### TROSY-HSQC NMR Spectra of *Aha* TRAP and dTRAP variants

Purified [U-^15^N]-TRAP samples were dialyzed against buffer A (100 mM NaCl, 50 mM NaPO_4_, pH 8.0, 0.02% NaN_3_) supplemented with 10% (v/v) D_2_O (99.8 atom %, Sigma-Aldrich) and 0.001% (v/v) DSS (4,4-Dimethyl-4-silapentane-1-sulfonic acid) (Sigma-Aldrich). 2D ^1^H-^15^N TROSY-HSQC NMR spectra of TRAP samples loaded in Wilmad^®^ 5 mm 541-PP NMR tubes (Sigma-Aldrich) were recorded at 55 °C (328K) on a Bruker Avance III HD Ultrashield 600 MHz NMR spectrometer equipped with a 5 mm triple-resonance inverse (TXI) cryoprobe with z-axis gradients. Spectra included: 500 µL of 62 µM TRAP (744 µM WT Trp binding sites), alone and titrated with 30 µL 11.5 mM Trp ([Trp]/[WT sites] ratio of 0.9). NMR spectra of dTRAP variants (WT-WT, WT-Mut, and Mut-WT) were recorded at 55 °C (328K) on a Bruker Avance III HD 800 MHz NMR spectrometer equipped with a TXI cryoprobe. The dTRAP NMR spectra were recorded on: a) 600 µL of 145.2 µM dTRAP WT-WT (1.74 mM WT Trp binding sites) alone and after addition of 77 µL of 14.6 mM Trp ([Trp]/[WT sites] ratio of 1.1); b) 555 µL 58 µM WT-Mut dTRAP (348 µM WT Trp binding sites) alone and after addition of 27 µL of 10.7 mM Trp stock ([Trp]/[WT sites] ratio of 1.5); c) 555 µL of 136 µM Mut-WT dTRAP (816 µM Trp WT binding sites) alone and after addition of 54.6 µL of 24 mM Trp ([Trp]/[WT sites] = 2.9).

### dTRAP Structure modelling and prediction

The starting model of dTRAP WT-WT with (G_4_S)_2_ linker and C-terminal TEV ENLYFQ was built using the loop modeling tools in Rosetta3 (RosettaRemodel release r231, 2019.35) [42,49] running on NMRBox [50], expanded via C6 symmetry and relaxed using built-in scoring functions. The lowest energy of 100 modeled trajectories was used as the starting point for modeling the cryoEM density. The position of the modeled linker extends from the C-terminus of the first protomer up through the center of the TRAP ring to the N-terminus of the second protomer, opposite of the Trp binding pocket and RNA binding interface.

An Alphafold2 model of dTRAP WT-WT was generated using Alphafold2 multimer v3 on Colabfold [51]. The hexameric dTRAP WT-WT sequences were aligned using MMseqs2 searching against PDB100 and in unpaired_paired mode. The top hit ranked by the predicted local distance difference test score (pLDDT) was relaxed using the Amber forcefield with the default settings. This model was qualitatively similar to that generated with Rosetta.

### RNA filter binding assay

To determine binding affinity of protein to RNA, nitrocellulose filter binding was performed in excess tryptophan (1 mM) with protein ring concentrations from 0 nM to 300 nM. A DNA sequence encoding the 55-nucleotide RNA, formed by 11 repeats of the sequence 5’-GAGUU-3’, was cloned into the plasmid pUC118GAGUU which includes a hammerhead ribozyme fused to the 3’-end [52]. The RNA (GAGUU)_11_ was generated from *in vitro* transcription followed by hammerhead ribozyme cleavage as previously described [53]. The RNA was 5’ end-labeled with and purified from 8 M urea gels by the crush and soak method, as described previously [52]. The reactions contained 5 pM RNA and were incubated at 37 °C for 10 minutes before filtering. The data were fit with a hyperbolic single binding site model, 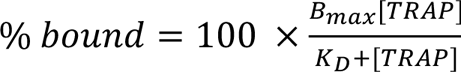, where *B*_max_ is the maximum observed bound fraction, [TRAP] is the total protein concentration, and *K*_D_ is the apparent equilibrium dissociation constant.

### Native mass spectrometry (nMS)

Trp-TRAP titrations monitored by nMS were performed at room temperature (∼25°C) with 5 µM protein sprayed from 400 mM AmAc on a Q Exactive Ultra-High Mass Range (UHMR) orbitrap mass spectrometer (Thermo Fisher Scientific). The instrument was modified to allow for surface induced dissociation, similar to a previously described modification.[16] A series of Trp-TRAP samples was prepared, each mixed with varying concentration of Trp: 0 µM, 10 µM, 20 µM, 30 µM, 40 µM, 50 µM, 60 µM, 70 µM, 80 µM, and 120 µM. Similarly, parallel samples of 2 µM dTRAP were prepared, mixed with Trp concentrations incremented by 2 µM from 0 µM to 24 µM, plus 48 µM. Double deconvolution of the m/z spectra to include proteoforms was applied to generate clean zero-charge mass spectra [54]. After deconvolution, both TRAP and dTRAP concentration correction was performed using mass balance (eq. 1), allowing the site concentration to be a fitted parameter, resulting in adjustment to 1.70 µM for dTRAP and 5.16 µM for *Bha* TRAP (Figure S 19). Next, the nMS data were globally fit with a nearest-neighbor (NN) model via *itcsimlib,* as previously described [16].

For the nMS titration of Trp into WT-WT dTRAP (2 µM ring),WT-Mut dTRAP (2 µM ring), Mut-WT dTRAP (2 µM ring), and Mut-Mut dTRAP (1 µM ring), experiments were conducted at room temperature (∼25°C) in 200 mM AmAc on an Exactive EMR Plus Orbitrap (Thermo Fisher Scientific) modified with a custom mass selection quadrupole and a surface-induced dissociation device [55]. The nMS titration of dTRAP WT-WT and Mut-Mut were performed with the protein samples mixed with Trp concentration of 0 µM, 12 µM, and 24 µM. The nMS titration of both dTRAP WT-Mut and dTRAP Mut-WT were performed with the protein samples mixed with Trp concentration of 0 µM, 36 µM, 64 µM, and 128 µM. The resulting data were deconvolved using MetaUniDec V4.4 [56]. The nMS data were also fit using the NN model as mentioned above.

### Cryo-EM sample preparation

An aliquot of 10 mg/mL of apo dTRAP was diluted with cryo-EM buffer (20 mM HEPES, 200 mM NaCl, pH 8) supplemented with a 0.6% stock of Triton X-100 to obtain a final sample of 6 mg/mL protein and 0.05% Triton X-100. Holo dTRAP was prepared by diluting an aliquot of 10 mg/mL protein with cryo-EM buffer supplemented with 1 mM Trp and 0.6% of Triton X-100, to obtain a sample at 6 mg/mL with 0.05% Triton X-100 and 600 μM Trp. Inclusion of Trion X-100 was found to be critical for avoiding adherence of TRAP rings to the air-water interface, and overcoming orientation bias [57]. The sample of Trp-bound WT-Mut dTRAP was prepared by diluting 14 mg/mL of WT-Mut dTRAP with 20 mM Trp in cryo-EM buffer and 0.6% Triton X-100 to obtain a final protein concentration of 6 mg/mL with 0.05% Triton X-100 and 1.4 mM Trp.

For all samples, Au Quantifoil R1.2/1.3 300 mesh grids were glow discharged (20 mA, 30 sec hold, 1 min glow discharge) using a PELCO easiGlow Discharge System. For all three samples, a volume of 3 μL was applied to the glow discharged grids and incubated horizontally for 60 seconds. Grids were then loaded onto a Vitrobot Mark IV and blotted with a blot force of 1 for 4 seconds at 4°C and 100% relative humidity before being plunge frozen into liquid ethane.

### Cryo-EM data collection

For each data set, movies were acquired using a total electron fluence of 60 e^-^/Å^2^ spread over 45 frames on a Thermo Fisher Scientific Krios G3i operating at 300 keV equipped with a Gatan K3 detector and Biocontinuum energy filter; slit widths were 15 eV for apo and holo dTRAP, and a 20eV for holo WT-Mut dTRAP. All movies were collected in super-resolution, counting mode using a nominal magnification of 81,000x (0.899 Å/px). For apo dTRAP, 3753 movies were collected over a defocus range of 0.5 - 0.2 μm. For holo dTRAP, 2307 movies were collected over a defocus range of 0.5 - 2.5 μm. For holo WT-Mut dTRAP, 1722 movies were collected over a defocus range of 1 – 2.4 μm.

### Cryo-EM data processing

For apo and holo dTRAP WT-WT, we performed data processing using CryoSPARC v2.15.0 [58]. Movies were imported and aligned using patch motion correction with an applied B-factor of 250, a maximum resolution of 3 Å, and subsequently binned by a factor of two to arrive at the physical pixel size of 0.899 Å/px. Patch contrast transfer function (CTF) estimation was performed with a maximum resolution of 2.5 Å. Manual curation of micrographs was used to remove micrographs with a CTF estimation > 5 Å and total motion > 100 Å.

For apo dTRAP, blob picking was used with a min/max particle diameter of 85/105 Å to process 100 micrographs. 2D classification was performed with 50 classes using a 130 Å mask to classify the good picks from noise/ice picks resulting in a selection of 15,612 particles to train a TOPAZ model^3^. TOPAZ was then used to particle pick 3,753 micrographs, resulting in 1.1 M picked particles. Particles were extracted with a box size of 256 px (0.899 Å/px) and binned to a pixel size of 3.596 Å. A single round of 2D classification was performed to remove 200k junk particle picks (overlapping particles, vitreous ice). Three ab-initio models were generated and subjected to multiple, iterative rounds of heterogenous refinement, homogeneous refinement, and 2D classification to remove particles that resulted in low resolution reconstructions. A final set of 91,038 particles were extracted with a box size of 256 pixels (0.899 Å/px), and subjected to homogenous refinement, giving way to a map with a resolution of 4.56 Å, and a subsequent non-uniform refinement that resulted in a map with a global resolution of 4.24 Å[59].

For holo dTRAP, blob picking was used with a min/max particle diameter of 90/110 Å to pick 30 micrographs. 2D classification was performed with 50 classes using a 140 Å mask to classify the good picks from noise/ice picks resulting in a selection of 7,849 particles that were used as templates to pick 2.3M “particles”. Three rounds of 2D classification were used to remove junk particle picks and produce a set of 612,271 particles that were then extracted with a box size of 256 px (0.899 Å/px) and binned to a pixel size of 3.596 Å. Four ab-initio models were generated and subjected to multiple rounds of heterogenous refinement, homogeneous refinement, and 2D classification to remove particles that resulted in low resolution reconstructions. A final set of 104,714 particles were extracted with a box size of 256 pixels (0.899 Å/px), subjected to homogenous refinement giving way to a map with a resolution of 4.30 Å, and a subsequent non-uniform refinement that resulted in a map with a global resolution of 3.6 Å[59]

For holo WT-Mut dTRAP, we performed data processing using CryoSPARC v3.2.0[58] . Movies were imported and aligned using patch motion correction with an applied B-factor of 500, a maximum alignment resolution of 5 Å. Patch CTF estimation was performed with a maximum resolution of 4 Å. Manual curation of micrographs was used to remove any micrographs that had a CTF resolution estimation > 5 Å and total motion > 100 Å. After curation, blob picking was used with a min/max particle diameter of 90/110 Å to pick 2.5M particles. 2D classifications was performed with 50 classes using a 140 Å mask to classify the good picks from noise/ice picks resulting in a selection of 402K particles that were used as template to pick 2.7M particle. Seven rounds of 2D classification were used to remove junk particle picks and produced a set of 182,939 particles that were then extracted with a box size of 512 pix and binned to a box size of 256 pix (0.899 Å/px). Three ab-initio models were generated and subjected to multiple rounds of heterogenous refinement and homogenous refinement followed by another three rounds of 2D classification to remove particles that resulted in low resolution reconstructions. A final set of 46,244 particles were extracted and subjected to multiple rounds of non-uniform refinement and local refinement generating a map with a resolution of 4.14 Å.

### Cryo-EM model building

For both apo and holo dTRAP, the Rosetta designed dTRAP model was used as a starting model. The C6 reconstructed map and initial model were used to refine a linked dTRAP protomer using multiple iterations of Coot, ISOLDE, and PHENIX Real Space Refinement [43–46]. Secondary structures were restrained during PHENIX real space refinement automatically using the HELIX/SHEET records form the crystal structure of *Aha* TRAP (PDB:3ZZL). Upon convergence of the model with the map, the map symmetry tool in Phenix was used to determine the NCS operators and to generate the other asymmetric units.

For Trp loaded WT-Mut dTRAP, the Cryo-EM structure of holo dTRAP was used as a starting model. The C6 symmetry reconstructed map and initial model were used to refine a single linked dimer using multiple iterations of ISOLDE and Phenix Real Space Refinement^6–8^. The last five rounds of iterations were performed using the full dTRAP ring constructed using the apply NCS operators’ tool by PHENIX so that the interface between the protomers is properly modeled.

## Abbreviations

nMS: native mass spectrometry
cryo-EM: cryogenic electron microscopy
NMR: nuclear magnetic resonance
Trp: tryptophan
TRAP: *rp* RNA-binding attenuation protein
*Aha*: *Alkalihalobacillus halodurans*
dTRAP: a protein construct composed of two *Aha* TRAP chains connected by a flexible linker
WT-WT/WT-Mut/Mut-WT/Mut-Mut dTRAP: dTRAP variants in which the first and second protomer are either the wild-type sequence (WT), or a Trp binding-defective mutant (Mut)
NN: nearest-neighbor
ITC: isothermal titration calorimetry.

## Acknowledgements

This work was supported by grants from the NIH to MPF, PG and VW (R01 GM077234, R01 GM062750, R01 GM120923, P41 GM128577). Chunhua Yuan and the Campus Chemical Instrument Center provided support for and access to NMR instrumentation. Electron microscopy data collection and analysis was performed at the Center for Electron Microscopy and Analysis (CEMAS) at The Ohio State University, with assistance from Yoshi Narui. Oliver Clarke (Columbia University), Rado Danev (University of Tokyo), and Alexis Rohou (Genentech) provided guidance the Cryo-EM data processing steps. Molecular graphics and analyses performed with UCSF ChimeraX, developed by the Resource for Biocomputing, Visualization, and Informatics at the University of California, San Francisco, was supported by NIH R01 GM129325 and the Office of Cyber Infrastructure and Computational Biology, NIAID. The work made use of NMRbox: National Center for Biomolecular NMR Data Processing and Analysis, a Biomedical Technology Research Resource (BTRR), which is supported by NIH grant P41 GM111135.

## Notes

### Competing Interest Statement

The authors have declared no competing interest.

