## Supporting Information for "Structural basis of nearest-neighbor cooperativity in the ring-shaped gene regulatory protein TRAP from protein engineering and cryo-EM"

A preprint of this article has been posted on bioRxiv under a CC-BY-NC-ND license.

Classification: Biological Sciences, Biophysics and Computational Biology

A

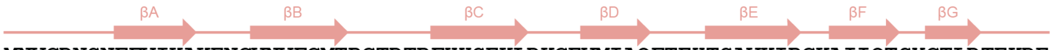


*Aha* TRAP MNVGDNSNFFVIKAKENGVNVFGMTRGTDTRFHHSEKLDKGEVMIAQFTEHTSAVKIRGKAIIQTSYGTLDTEKDE

1 10 20 30 40 50 60 70

B

### Expected protein sequence

TRAP WT

*Expression, expected protomer mass: 10167.28 Da*

**MNVGDNSNFFVIKAKENGVNVFGMTRGTDTRFHHSEKLDKGEVMIAQFTEHTSAVKIRGKAIIQTSYGTLDTEKDEENLYFQGHHHHHH**

*Post-purification, expected protomer mass: 9287.38 Da*

**MNVGDNSNFFVIKAKENGVNVFGMTRGTDTRFHHSEKLDKGEVMIAQFTEHTSAVKIRGKAIIQTSYGTLDTEKDEENLYFQ**

C

**dTRAP WT-WT**

*Post-purification, expected protomer mass: 18392.45 Da*

**MNVGDNSNFFVIKAKENGVNVFGMTRGTDTRFHHSEKLDKGEVMIAQFTEHTSAVKIRGKAIIQTSYGTLDT
EKDEGGGGSGGGGSMNVGDNSNFFVIKAKENGVNVFGMTRGTDTRFHHSEKLDKGEVMIAQFTEHTSAVKIR
GKAIIQTSYGTLDTEKDEENLYFQ**

**dTRAP Mut-WT**

*Post-purification, expected protomer mass: 18332.40 Da*

**MNVGDNSNFFVIKAKENGVNVFGMTRGTDTRFHHSEKLDKGEVMIAQFAEHASAVKIRGKAIIQTSYGTLDT
EKDEGGGGSGGGGSMNVGDNSNFFVIKAKENGVNVFGMTRGTDTRFHHSEKLDKGEVMIAQFTEHTSAVKIR
GKAIIQTSYGTLDTEKDEENLYFQ**

**dTRAP WT-Mut**

*Post-purification, expected protomer mass: 18332.40 Da*

**MNVGDNSNFFVIKAKENGVNVFGMTRGTDTRFHHSEKLDKGEVMIAQFTEHTSAVKIRGKAIIQTSYGTLDT
EKDEGGGGSGGGGSMNVGDNSNFFVIKAKENGVNVFGMTRGTDTRFHHSEKLDKGEVMIAQFAEHASAVKIR
GKAIIQTSYGTLDTEKDEENLYFQ**

**dTRAP Mut-Mut**

*Post-purification, expected protomer mass: 18272.35 Da*

**MNVGDNSNFFVIKAKENGVNVFGMTRGTDTRFHHSEKLDKGEVMIAQFAEHASAVKIRGKAIIQTSYGTLDT
EKDEGGGGSGGGGSMNVGDNSNFFVIKAKENGVNVFGMTRGTDTRFHHSEKLDKGEVMIAQFAEHASAVKIR
GKAIIQTSYGTLDTEKDEENLYFQ**

D

### Expected RNA sequence

*$(GAGUU)_{11}$*

*expected mass: 18,223.97 Da*

**GAGUUGAGUUGAGUUGAGUUGAGUUGAGUUGAGUUGAGUUGAGUUGAGUUGAGUU**

1 3 6 9 11

*Figure S 1. Sequence information of monomeric Aha TRAP, dimeric Aha dTRAP and single stranded RNA $(GAGUU)_{11}$.*

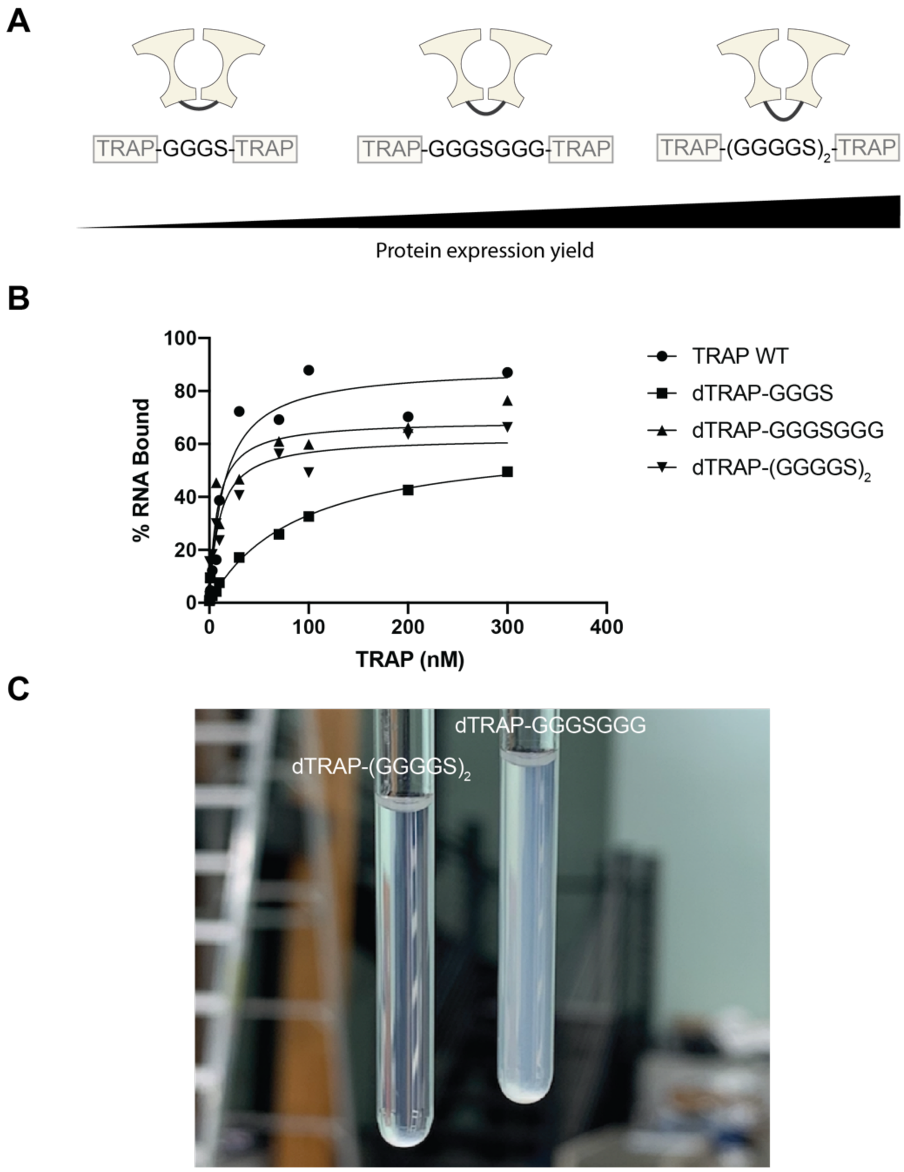


*Figure S 2. Aha dTRAP construct with the linker of $(GGGGS)_2$ was selected for experiments. (A) Schematic of dTRAP with different Gly/Ser linkers. dTRAP constructs were ranked by the order of protein expression yield in E. coli BL21(DE3). The yield of dTRAP-GGGS is extremely low. (B) Binding affinity of dTRAP constructs to RNA was determined using RNA filter binding assay in the presence of excess Trp. Lines fit to the data points demonstrate the nonlinear single binding site equation fit. Both dTRAP-$(GGGGS)_2$ ($K_d$ = 7 nM) and dTRAP-GGGSGGG ($K_d$ = 10 nM) share the similar binding affinity with TRAP WT ($K_d$ = 5 nM). But the RNA binding affinity of dTRAP-GGGS ($K_d$ = 90 nM) drops more than 10 times, showing defects of RNA binding in dTRAP-GGGS design. (C) A photo of purified dTRAP proteins in NMR tubes. After purification and refolding, dTRAP-GGGSGGG tended to precipitate as the solution went milky, whereas dTRAP-$(GGGGS)_2$ was more soluble and stable.*

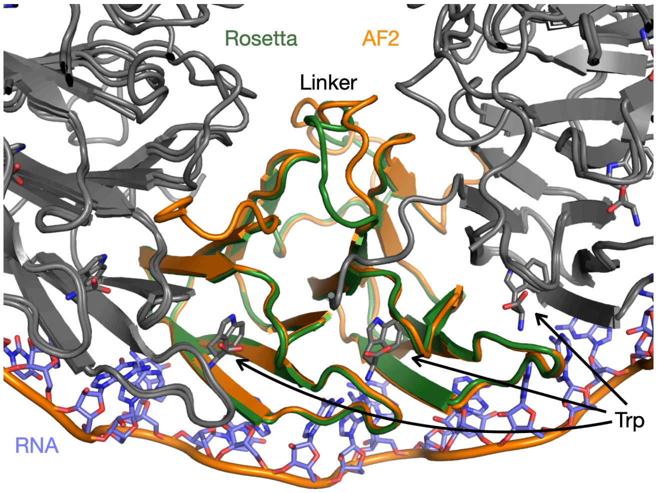


*Figure S 3.* Inter-*protomer linker is not predicted to interfere with Trp or RNA binding. Comparisons of Rosetta (Green) and AlphaFold2 (Orange) modelled structures of Aha dTRAP with a 10-residue linker (GGGGS)$_2$. RNA is modeled by homology to Gst TRAP bound to Trp and RNA (1C9S), by superposing in PyMOL chain Q on one chain of the Rosetta model.*

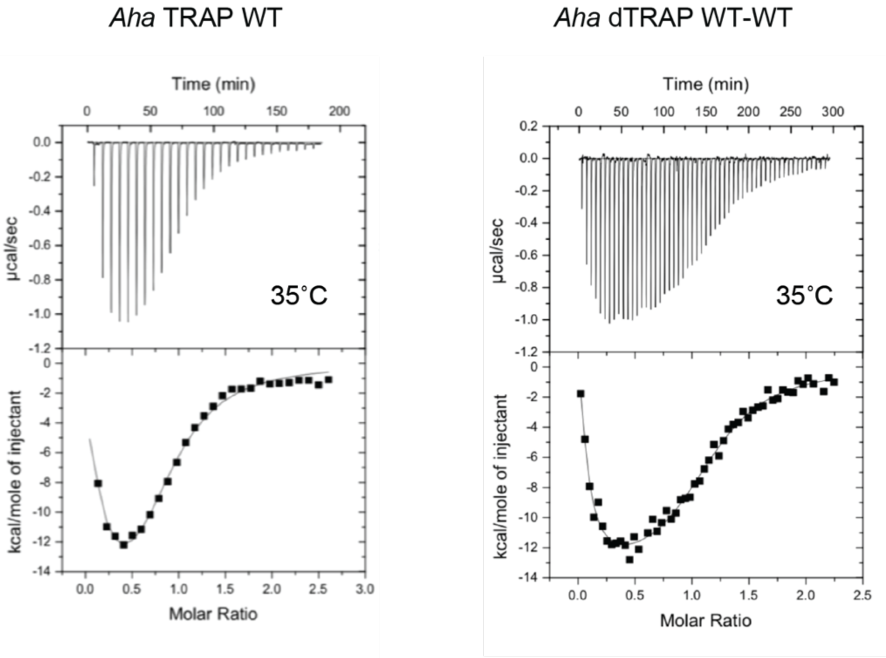


*Figure S 4. ITC thermograms of TRAP WT and dTRAP WT-WT present similar bi-phasic trends. The Trp- TRAP titrations were performed at 35˚C.*

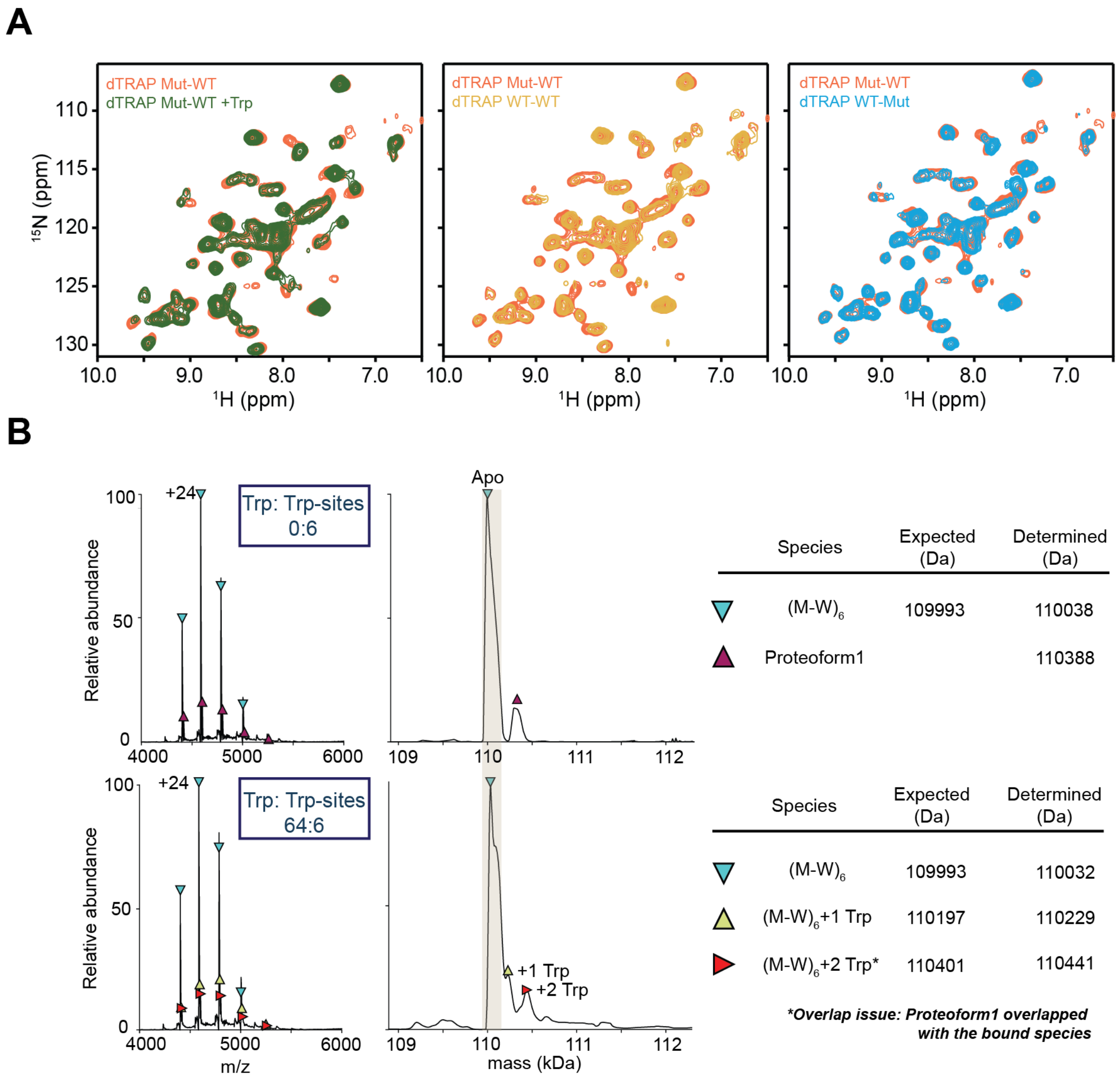


*Figure S 5. dTRAP Mut-WT is defective in Trp binding. (1) Comparative 2D $^{1}H$-$^{15}N$-correlated NMR spectra of Mut-WT dTRAP in the absence and presence of Trp, and the comparisons with WT-WT and WT-Mut dTRAP. Left: The chemical shifts of dTRAP Mut-WT in the presence of 0 Trp (Orange) and 8-fold excess Trp (Green) overlapped well, indicating defective Trp binding. Middle and Right: The peaks of dTRAP Mut-WT (Orange) superimposed well with dTRAP WT-WT (Yellow) and dTRAP WT-Mut (Blue) in the absence of Trp, indicating these three proteins share the similar structures. (2) Native mass spectra of dTRAP Mut-WT (2 µM) titrated against 0 and 64 µM Trp, respectively, showing that Mut-WT lack of ability to bind Trp.*

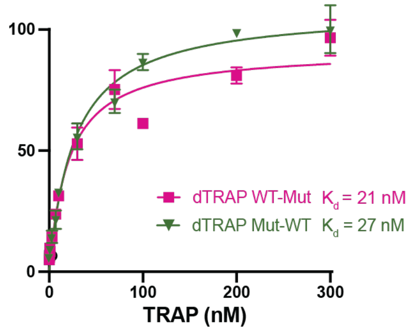


*Figure S 6. RNA filter binding assay, that was performed at the same condition as shown in Figure 4B, showed that dTRAP Mut-WT binds (GAGUU)$_{11}$ tightly in the presence of excess Trp.*

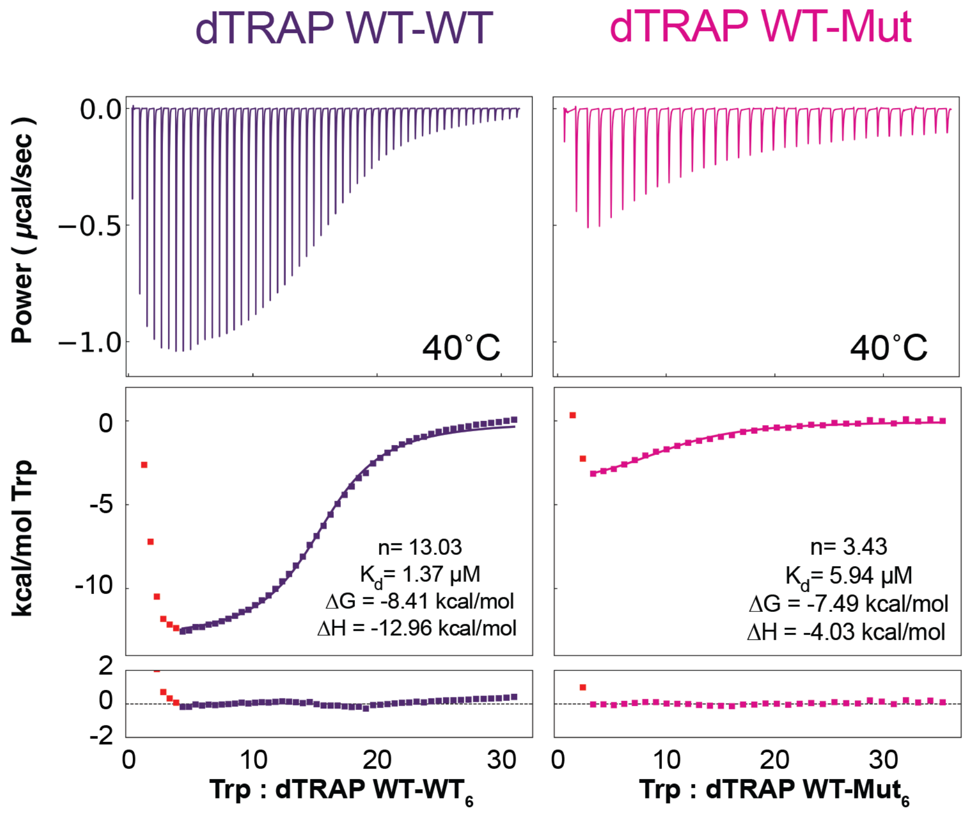


*Figure S 7. One-site model fitting of the second phase of ITC thermograms of Trp to dTRAP variants titrations; apparent model parameters as shown. The red points were omitted for fitting with this model.*

**A**

### *Bha* dTRAP WT-WT

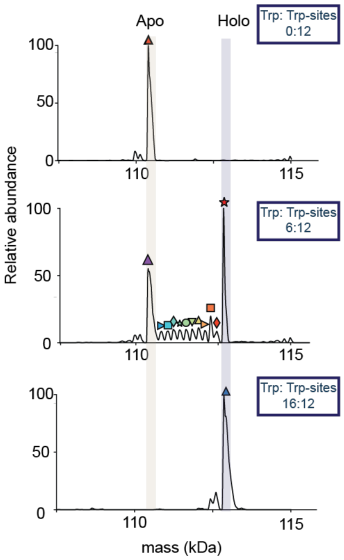


| Species | Expected (Da) | Determined (Da) |
|---|---|---|
| $(W\text{-}W)_6$ | 110355 | 110394 |
| $(W\text{-}W)_6$ | 110355 | 110398 |
| $(W\text{-}W)_6$+ 2 Trp | 110763 | 110244 |
| $(W\text{-}W)_6$+ 3 Trp | 111967 | 110468 |
| $(W\text{-}W)_6$+ 4 Trp | 111172 | 110664 |
| $(W\text{-}W)_6$+ 5 Trp | 111376 | 110876 |
| $(W\text{-}W)_6$+ 6 Trp | 111580 | 111102 |
| $(W\text{-}W)_6$+ 7 Trp | 111784 | 111323 |
| $(W\text{-}W)_6$+ 8 Trp | 111989 | 110468 |
| $(W\text{-}W)_6$+ 9 Trp | 112192 | 110664 |
| $(W\text{-}W)_6$+ 10 Trp | 112397 | 110876 |
| $(W\text{-}W)_6$+ 11 Trp | 112601 | 111102 |
| $(W\text{-}W)_6$+ 12 Trp | 112805 | 112832 |
| $(W\text{-}W)_6$+12 Trp | 112805 | 112861 |

**B**

### *Bha* dTRAP WT-Mut

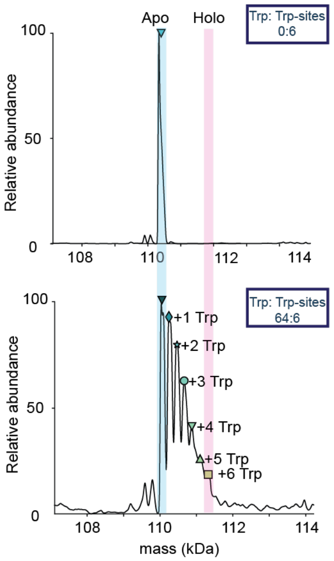


| Species | Expected (Da) | Determined (Da) |
|---|---|---|
| $(W\text{-}M)_6$ | 109993 | 110029 |
| $(W\text{-}M)_6$ | 109993 | 110036 |
| $(W\text{-}M)_6$+ 1 Trp | 110197 | 110244 |
| $(W\text{-}M)_6$+ 2 Trp | 110401 | 110468 |
| $(W\text{-}M)_6$+ 3 Trp | 110605 | 110664 |
| $(W\text{-}M)_6$+ 4 Trp | 110910 | 110876 |
| $(W\text{-}M)_6$+ 5 Trp | 111014 | 111102 |
| $(W\text{-}M)_6$+ 6 Trp | 111218 | 111323 |

*Figure S 8. Mass annotations of native MS titration spectra of dTRAP WT-WT and WT-Mut.*

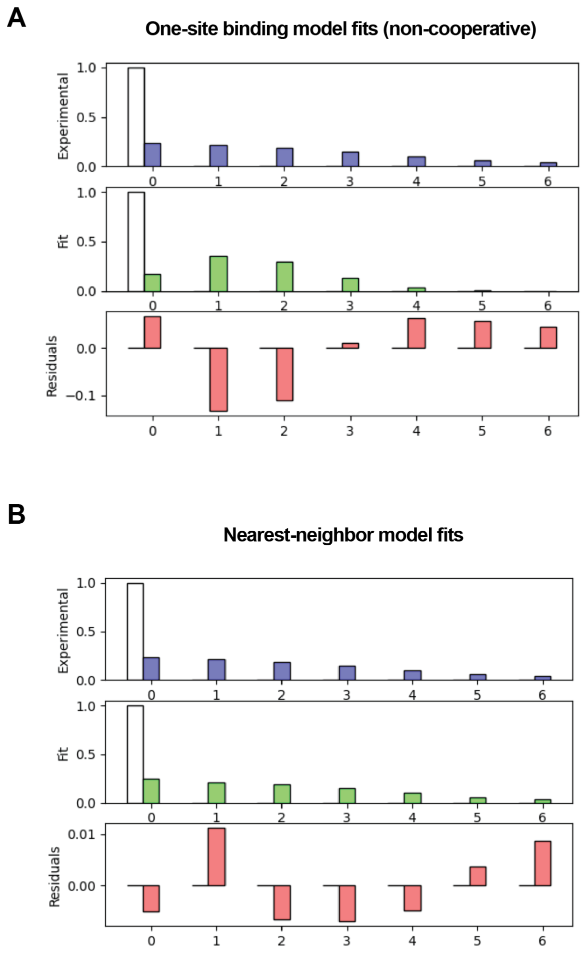


*Figure S 9. Trp to WT-Mut dTRAP titration monitored by native MS titration is not well explained by an independent sites model. Top, histogram of experimental and best fit populations with 0-6 bound Trp, and residuals, for an oligomer with six independent sites. Bottom, fits to a NN model. Populations from the data in (Figure 4D).*

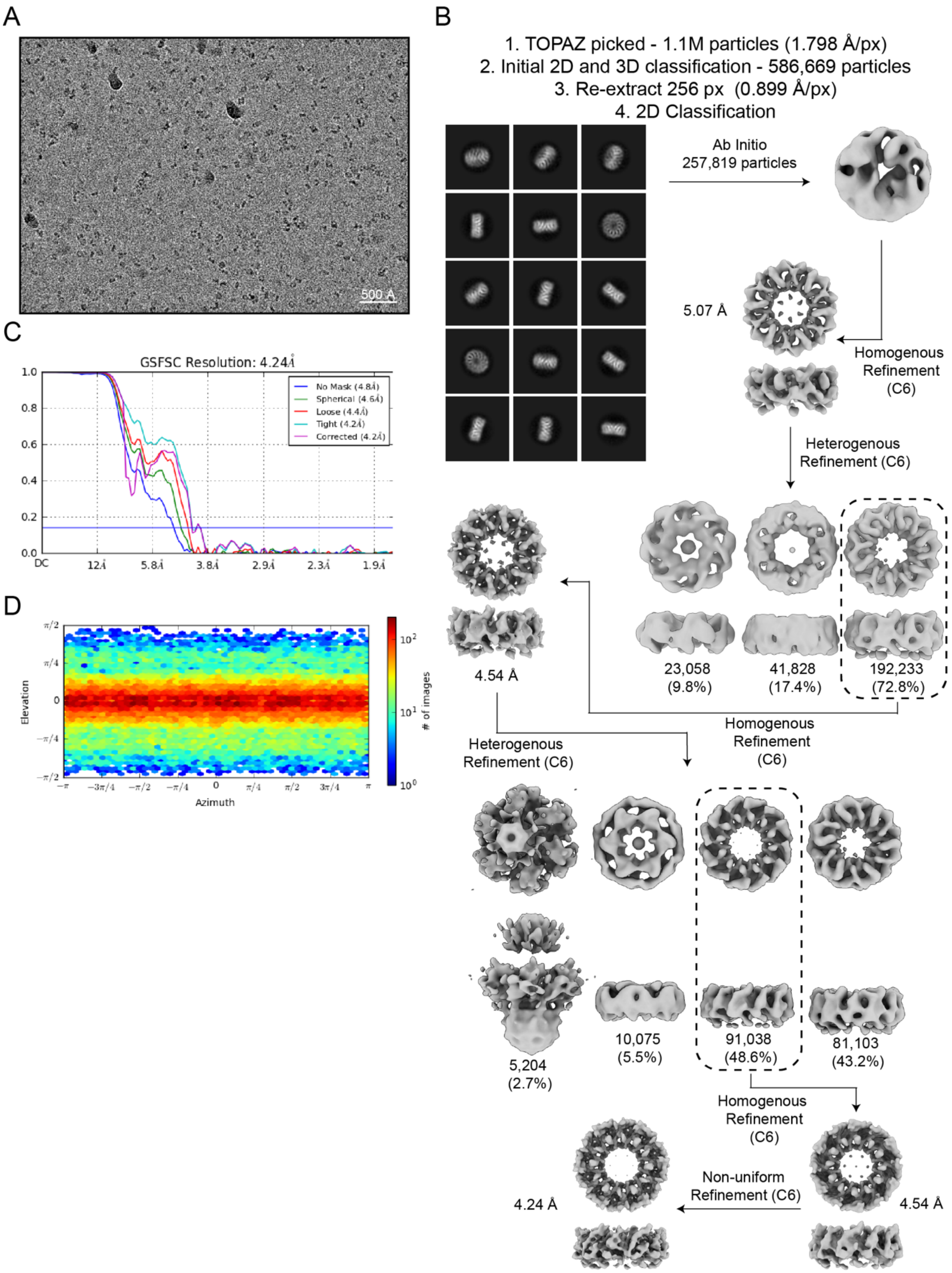


*Figure S 10. Cryo-EM data processing of apo WT Bha dTRAP. A. Representative micrograph. B. CryoSPARC classification and reconstruction workflow. Classes with dashed outline were selected for subsequent steps. C. Gold standard FSC resolution reported by CryoSPARC. D. Orientation estimation of the particle collected for final reconstruction.*

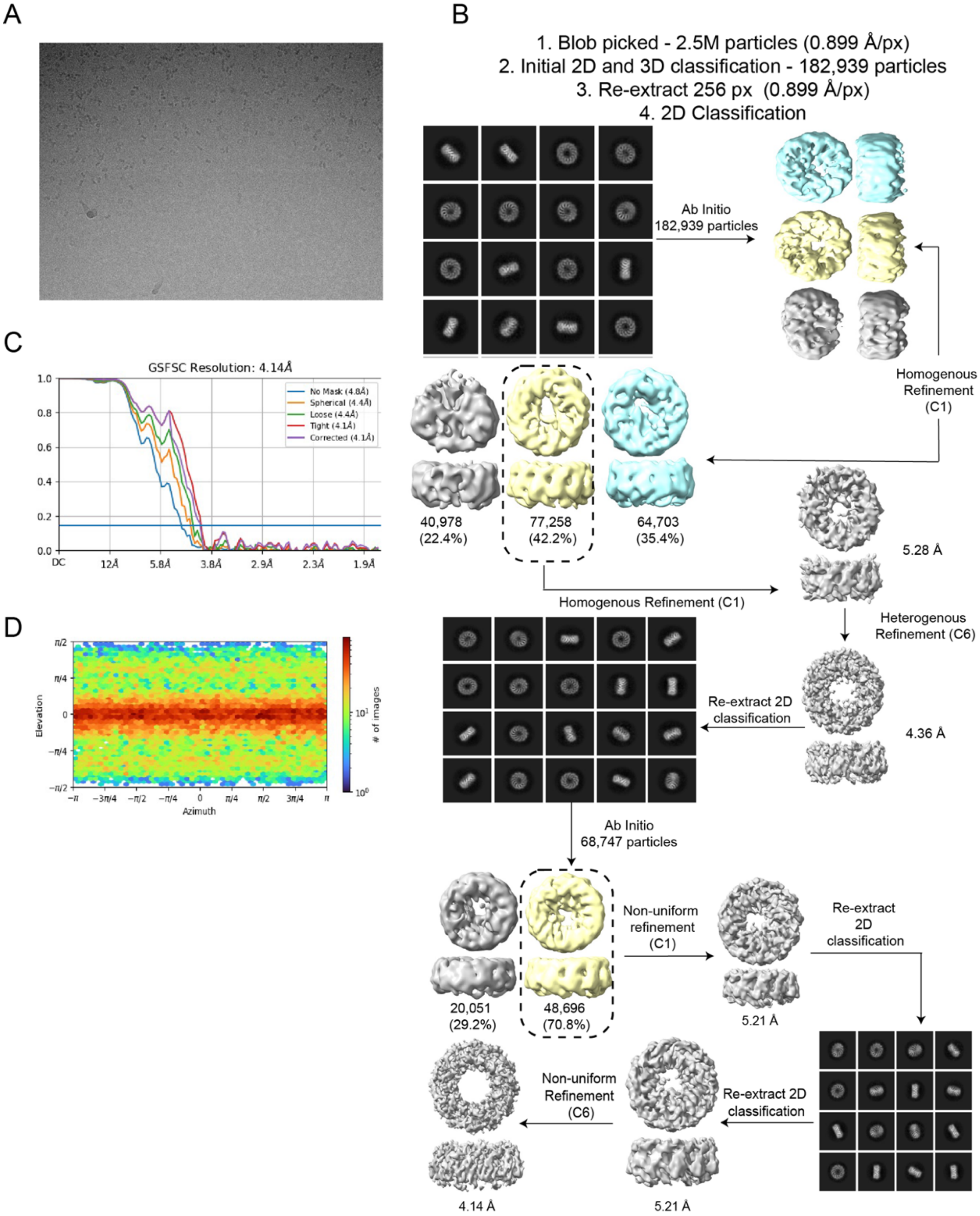


*Figure S 11. Cryo-EM data processing of holo WT-Mut Bha dTRAP. A. Representative micrograph. B. CryoSPARC classification and reconstruction workflow. Classes with dashed outline were selected for subsequent steps. C. Gold standard FSC resolution reported by CryoSPARC. D. Orientation estimation of the particle collected for final reconstruction.*

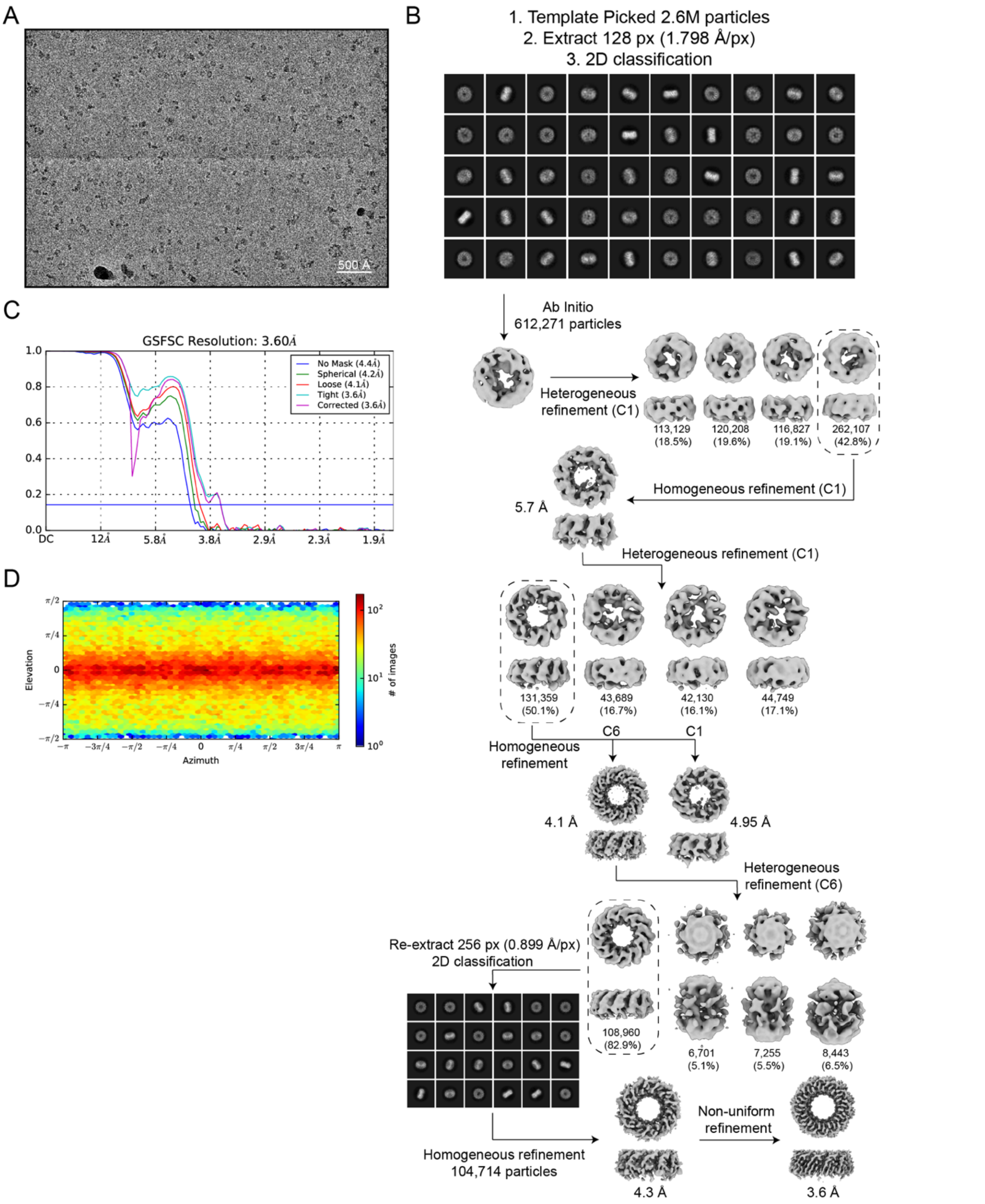


*Figure S 12. Cryo-EM data processing of holo WT Bha dTRAP. A. Representative micrograph. B. CryoSPARC classification and reconstruction workflow. Classes with dashed outline were selected for subsequent steps. C. Gold standard FSC resolution reported by CryoSPARC. D. Orientation estimation of the particle collected for final reconstruction.*

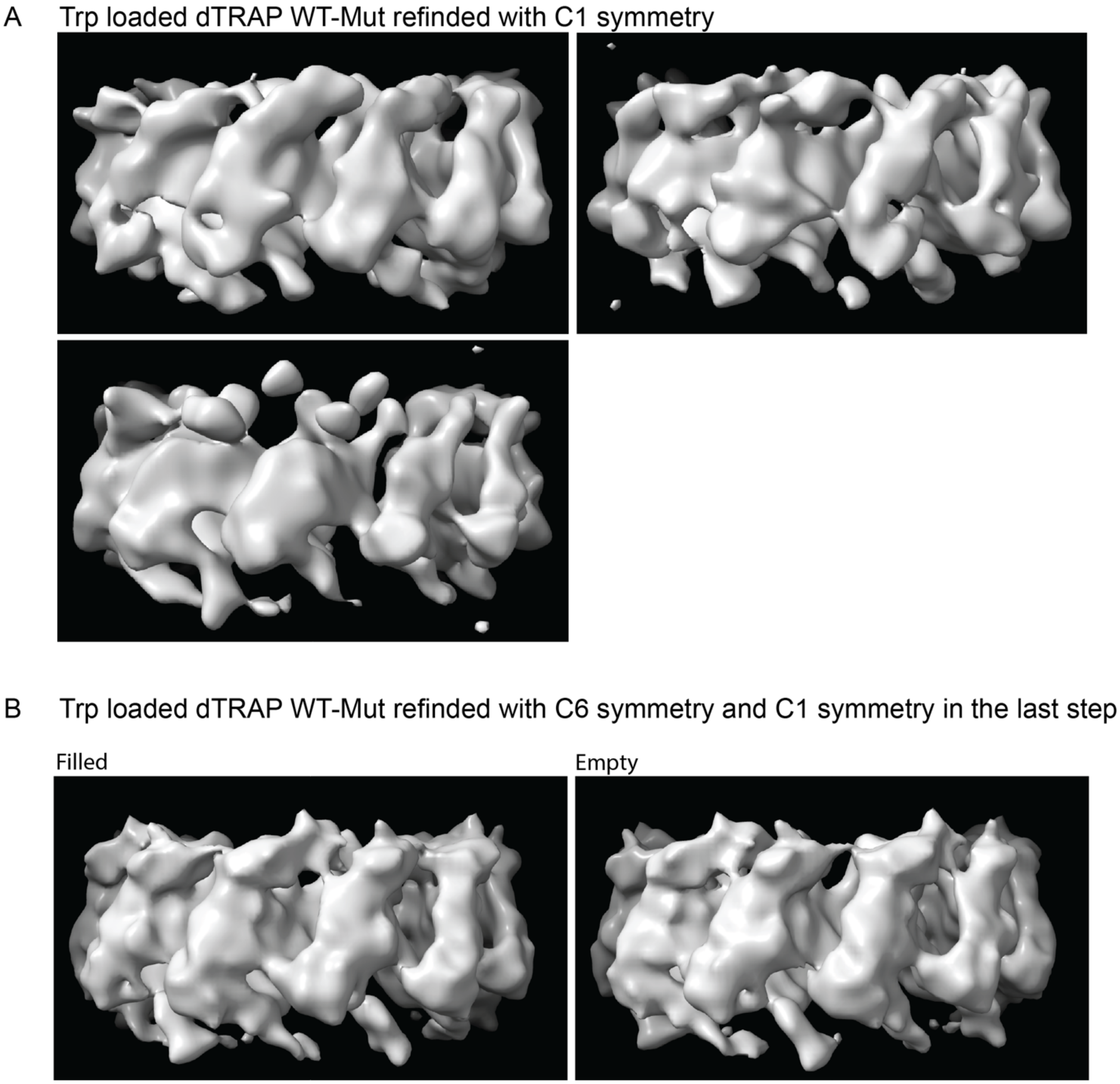


*Figure S 13. Cryo-EM map of Trp loaded dTRAP WT-Mut refined with different symmetry. A. Cryo-EM map refined using the same data set as Trp loaded dTRAP WT-Mut in figure 4 and 5. Workflow is show in Figure S13. This map is refined without application of symmetry (C1); contoured at a level of 0.0967 in ChimeraX. B. Map refined with C6 symmetry and then C1 symmetry in the last set using the same data set as A. Workflow is showing in figure S14. Showing with a contour level of 0.0645 in ChimeraX.*

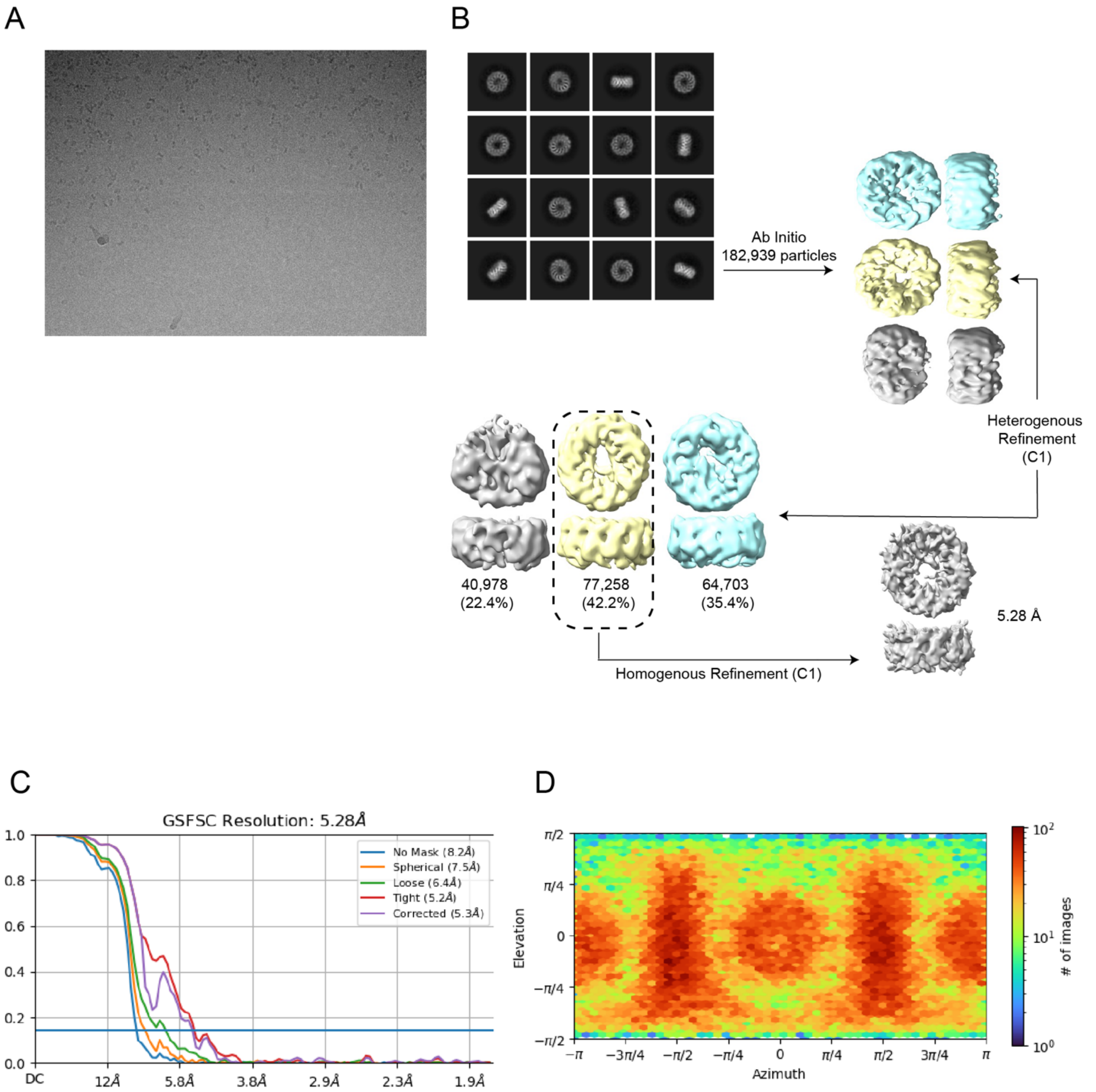


*Figure S 14. Cryo-EM map of Trp loaded dTRAP WT-Mut refined with only C1 symmetry using the same data set as Figure S 11. A. Representative micrograph. B. CryoSPARC classification and reconstruction workflow. Classes with dashed outline were selected for subsequent steps. C. Gold standard FSC resolution reported by CryoSPARC. D. Orientation estimation of the particle collected for final reconstruction.*

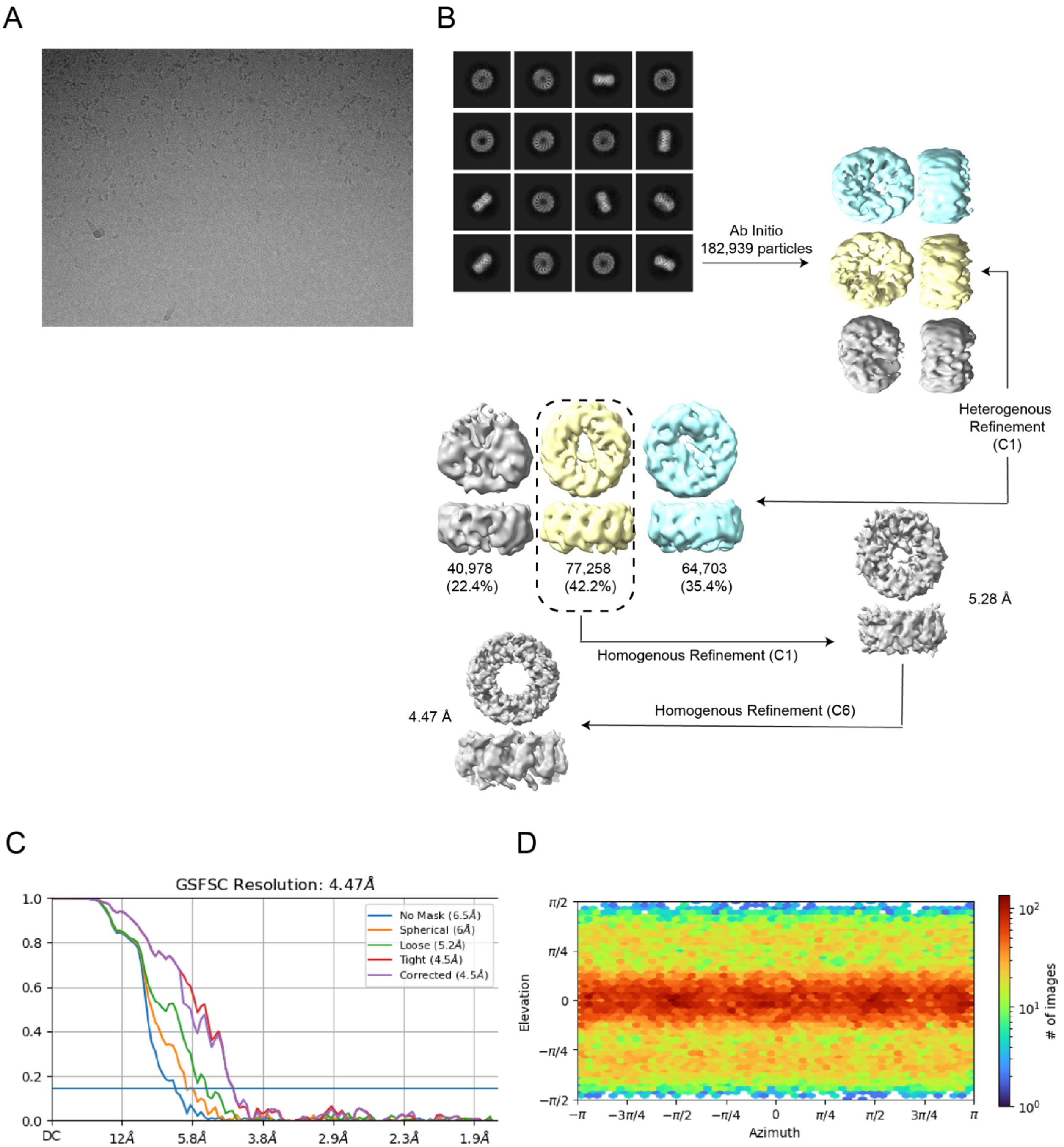


*Figure S 15. Cryo-EM map of Trp loaded dTRAP WT-Mut refined with C1 symmetry using the same data set as Figure S 11 and then C6 symmetry. A. Representative micrograph. B. CryoSPARC classification and reconstruction workflow. Classes with dashed outline were selected for subsequent steps. C. Gold standard FSC resolution reported by CryoSPARC. D. Orientation estimation of the particle collected for final reconstruction.*

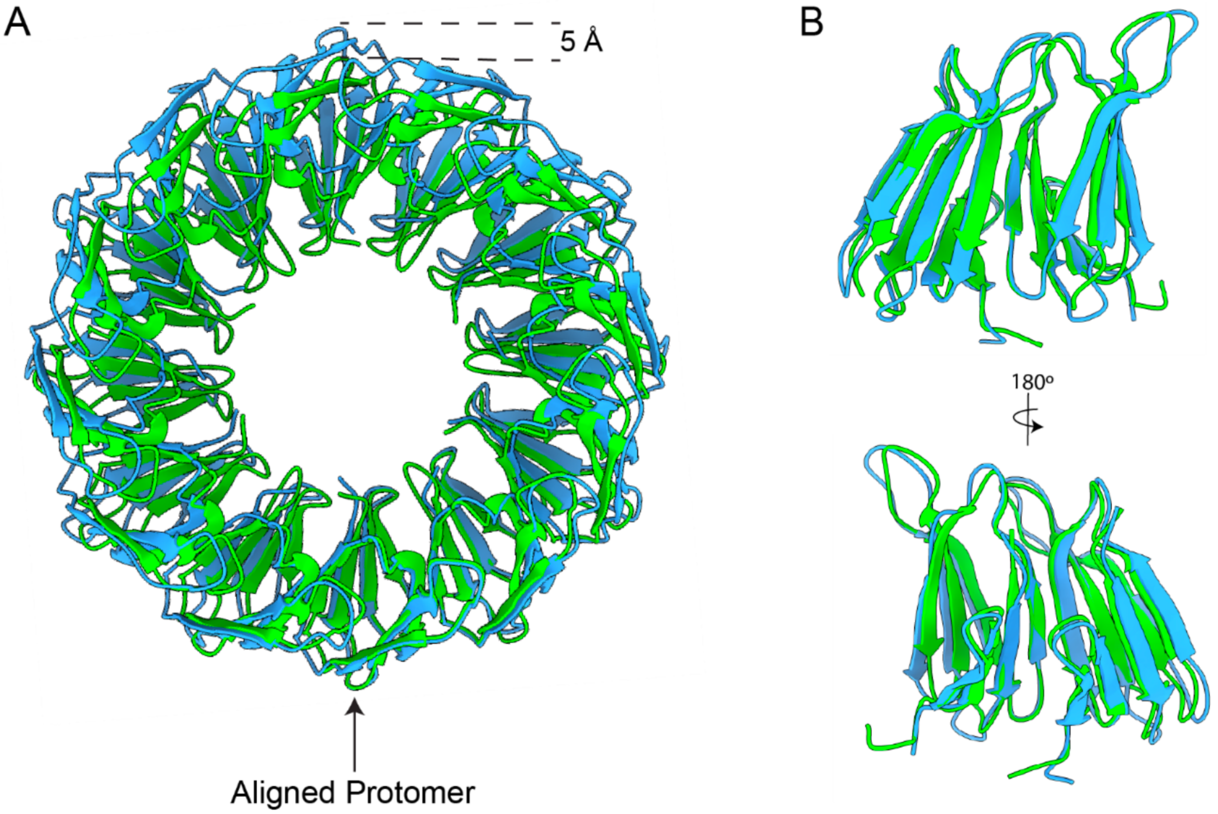


*Figure S 16. Apo and holo dTRAP rings are expanded by ~6% compared to WT TRAP rings. (A) Overlay of Aha WT TRAP (PDB 3ZZL, green) and the refined model of holo dTRAP WT-WT (blue). Chains were aligned on chain A of each model. Measurements taken between common Cα atoms. (B) Overlay of two Aha WT TRAP monomers with one holo dTRAP WT-WT dimer.*

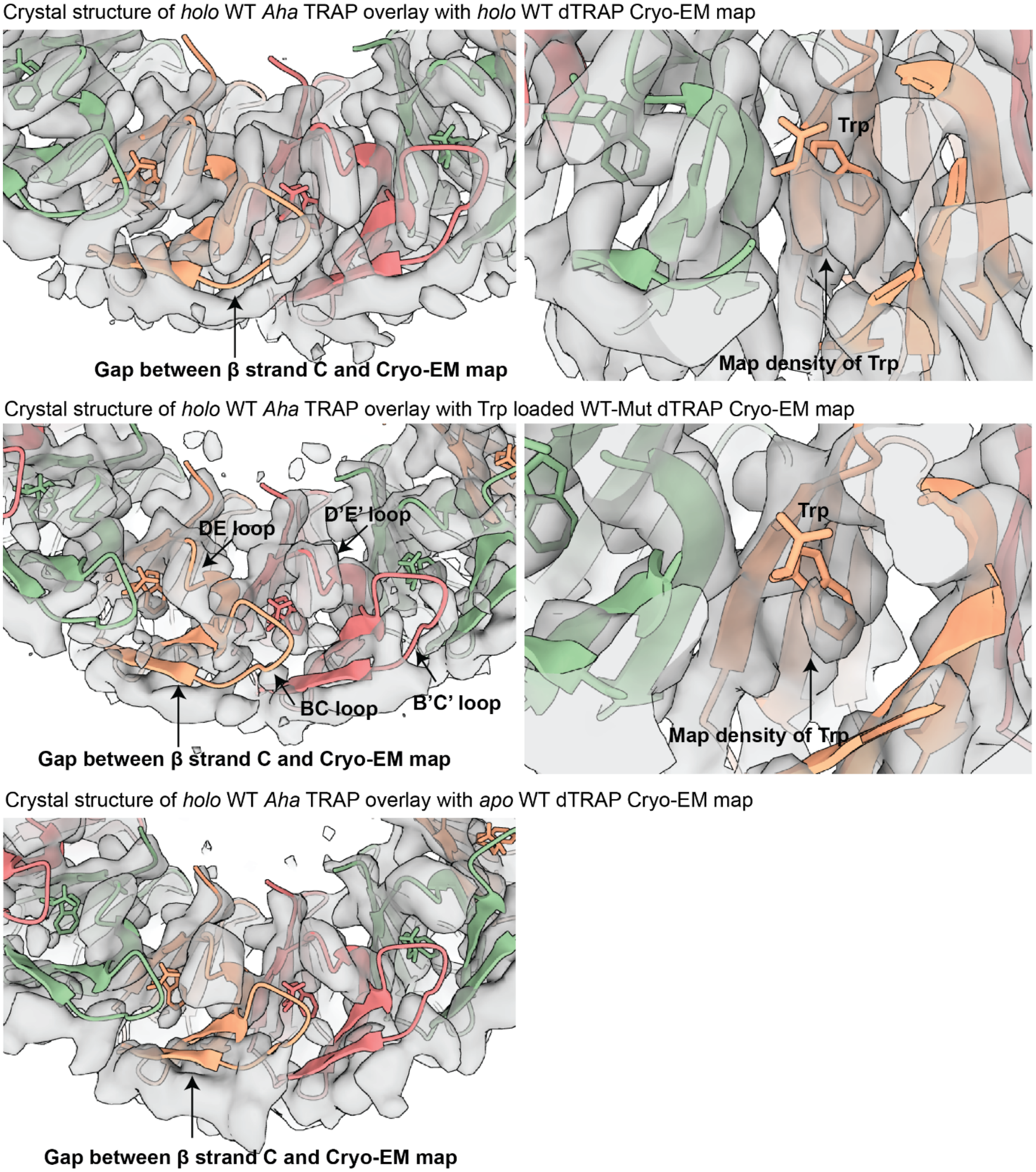


*Figure S 17. The crystal structure of Aha TRAP (3ZZL) does not fit the EM maps. Top panel: the crystal structure of Aha TRAP fitted into the CryoEM map of holo WT dTRAP (contour level set at 0.184 in ChimeraX). A obvious gap is found between the β stand C of the structure and the β stand C region of the map (left) suggesting the ring expansion of dTRAP. Similar conclusion can be drawn from comparing the position of Trp in the crystal structure to the Trp density observed in the EM map. The same ring expansion is also observed in the WT-Mut dTRAP map (middle, contour level set at 0.11 in ChimeraX), and apo dTRAP map (bottom, contour level set at 0.135 in ChimeraX).*

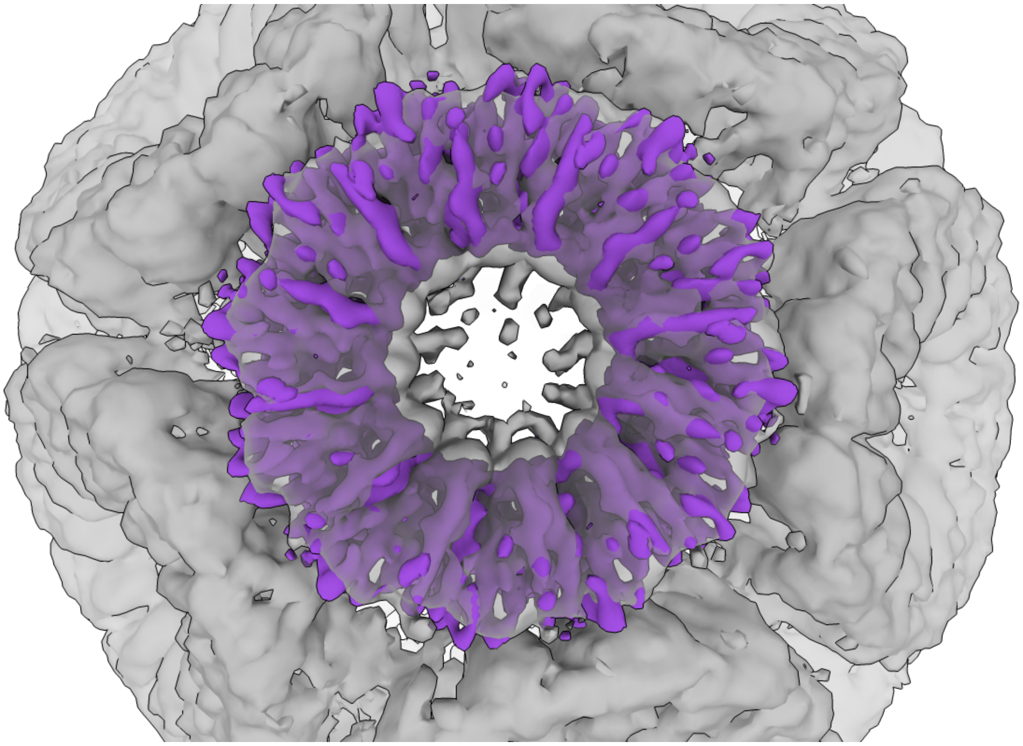

*Figure S 18. Density map for dTRAP is expanded relative to WT Aha TRAP. Map from Aha TRAP nano-cages is grey [18] (contour level 0.418 in ChimeraX). The holo dTRAP map is in purple (contour level 0.184 in ChimeraX. Maps were manually superposed in ChimeraX.*

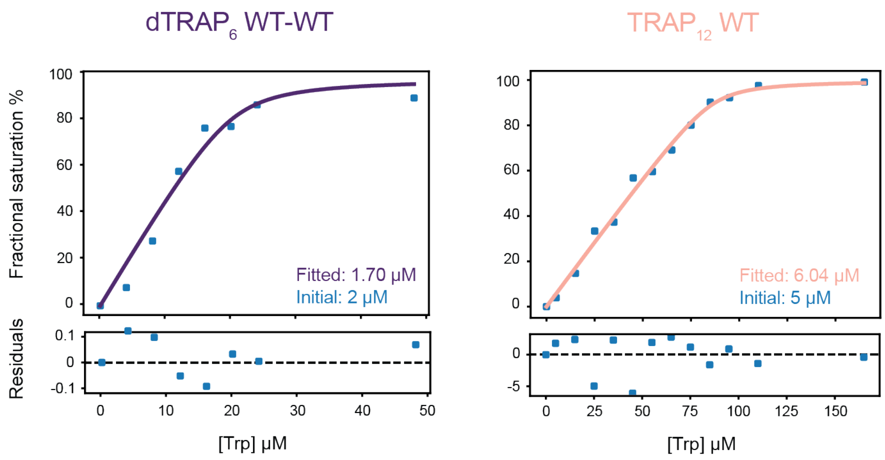


*Figure S 19. Representative concentration corrections of native MS data using mass balance by allowing site concentration to be an optimized parameter.*

*Table S1. CryoEM data collection parameters and structure validation report.*

| **Data Collection Parameters** | ***Aha* dTRAP *apo*** | ***Aha* dTRAP *holo*** | ***Aha* dTRAP WT-Mut** |
|---|---|---|---|
| Microscope | Titan Krios G3i | Titan Krios G3i | Titan Krios G3i |
| Voltage | 300 kV | 300 kV | 300 kV |
| Camera | Gatan K3 | Gatan K3 | Gatan K3 |
| Automation software | EPU | EPU | EPU |
| Number of collected micrographs | 3,753 | 2,307 | 1,721 |
| Nominal magnification | 81,000 | 81,000 | 81,000 |
| Pixel size | 0.889 Å (0.4495 Å super-res.) | 0.889 Å (0.4495 Å super-res.) | 0.899 Å (0.4495 Å super-res.) |
| Defocus range | 0.5 - 2.0 µm | 0.5 - 2.5 µm | 1.0-2.4 µm |
| Mean defocus | 1.25 µm | 1.5 µm | 1.7 µm |
| Electron fluence per frame | 1.33 e-/Å2 | 1.33 e-/Å2 | 1 e-/Å2 |
| Total electron fluence | 60 e-/Å2 | 60 e-/Å2 | 60 e-/Å2 |
| Number of frames per exposure | 45 | 45 | 60 |
| Illuminated area | 1.5 µm | 1.5 µm | 1.25 µm |
| C2 aperture | 50 µm | 50 µm | 50 µm |
| Objective aperture | 100 µm | 100 µm | 100 µm |
| Energy filter slit width | 15 eV | 15 eV | 20 eV |
| Grid type | Quantifoil Au R1.2/1.3 300 mesh | Quantifoil Au R1.2/1.3 300 mesh | Quantifoil Cu R 1.2/1.3 300 mesh |
| Specimen temperature | ≈86 K | ≈86 K | ≈86 K |
| **3D-Reconstruction Summary** | | | |
| Number of micrographs used | 3,545 | 1,956 | 1,721 |
| Number of particles picked | 586,669 | 612,271 | 182,939 |
| Number of particles in final reconstruction | 91,038 | 104,714 | 46,244 |
| Particle box size | 256 px | 256 px | 512 px |
| Resolution (0.143 FSC, masked) | 4.2 Å | 3.61 Å | 4.14 Å |

| | | | |
|---|---|---|---|
| Guinier B-factor | 208.9 Å$^2$ | 216 Å$^2$ | 247.6 Å$^2$ |
| **Model Composition** | | | |
| Number of asymmetric units | 6 | 6 | 6 |
| Non-hydrogen atoms | 11,676 | 13,056 | 12846 |
| Protein residues | 738 | 822 | 816 |
| **Refinement** | | | |
| Refinement software | Coot, ISOLDE, Phenix | Coot, ISOLDE, Phenix | ISOLDE, Phenix |
| FSC (map vs. refined model at 0.5/0.143) | 4.5/4.2 Å | 4.2/3.8 Å | 4.5/4.1 Å |
| Average cross correlation coefficient | 0.73 | 0.73 | 0.66 |
| Protein mean atomic B factor | 182.28 | 115.18 | 115.57 |
| **RMS Deviations** | | | |
| Bond Length (# > 4σ) | 0.003 Å (0) | 0.004 Å (0) | 0.004 Å (0) |
| Bond Angle (# > 4σ) | 0.912 ° (0) | 0.923 ° (0) | 0.942 ° (0) |
| **Validation** | | | |
| Molprobity Score | 1.30 | 1.58 | 1.57 |
| Clash score (all atoms) | 5.58 | 4.91 | 11.31 |
| $C_\beta$ outliers | 0% | 0% | 0% |
| CaBLAM outliers | 3.97% | 1.57% | 1.57% |
| **Ramachandran** | | | |
| Favored | 99.13% | 95.42% | 100% |
| Allowed | 0.87% | 4.58% | 0.00% |
| Outliers | 0% | 0% | 0.00% |
| Sheet Z-Score | -0.39 | 1.97 | 1.43 |
| Loop Z-Score | -2.32 | -1.58 | -1.47 |
